# Physical drivers of benthic and pelagic algal biomass dynamics in lakes: a conceptual exploration with ‘Lake2D’

**DOI:** 10.1101/2025.08.22.671802

**Authors:** Hugo Harlin, Karl Larsson, Åke Brännström, Sebastian Diehl

## Abstract

Size, depth and basin shape are important factors controlling the physics, chemistry and, ultimately, the productivity of lakes. To our knowledge, a comprehensive theoretical investigation of the physical determinants of lake primary production from a conceptual, process-based perspective has not been performed. To address this knowledge gap, we developed and analyzed ‘Lake2D’, a process-based, reaction-advection-diffusion model that adopts a 2-dimensional modeling approach by reducing lake bathymetry to its hypsographic depth distribution under the simplifying assumption of radial symmetry of the lake basin. In simpler terms, the model assumes the lake basin is perfectly circular and uses a cross-section to represent its depth and shape. We used Lake2D to explore and analyze the dynamics of pelagic and benthic algae on a whole-lake scale in response to six environmental drivers (lake area and depth, horizontal and vertical turbulent mixing, water transparency and nutrient status), which we varied over ranges that are representative of the vast majority of the world’s lakes. Numerical analyses reveal three distinct patterns across environmental parameter space. (1) Benthic algae dominate total biomass in shallow, clear lakes with low nutrient content. In these lakes, low horizontal and vertical mixing is beneficial to benthic but detrimental to pelagic algae, which experience high sinking losses, thus liberating nutrients and minimizing shading of benthic algae. (2) Increasing mean lake depth, abiotic turbidity and/or nutrient content strongly benefits pelagic algae, which increasingly shade out benthic algae and dominate total biomass. In these lakes, increased mixing affects both algal types positively due to increased nutrient transport to shallow, well-lit areas. (3) Finally, at high lake mean depth and/or abiotic turbidity, a large fraction of the lake volume is aphotic. In such lakes, high horizontal and vertical mixing is detrimental to pelagic algae but beneficial to benthic algae, because pelagic algae are mixed to aphotic depths, liberating nutrients and minimizing shading. Moving from shallow, clear lakes via deeper and/or nutrient-rich lakes to very deep and/or turbid lakes, benthic and pelagic algae therefore show opposite gradual biomass shifts in response to vertical and horizontal mixing. Specifically, benthic biomass is highest at low and lowest at high overall mixing in shallow, nutrient-poor lakes, but shows the opposite trend in all other lakes. Conversely, pelagic biomass is highest at low and lowest at high overall mixing in very deep and/or turbid lakes, but shows the opposite trend in all other lakes.

## 1 Introduction

Size, depth and basin shape are important factors controlling the physics, chemistry and, ultimately, the productivity of lakes. For example, deeper lakes are typically less completely mixed, are clearer, and have lower pelagic nutrient and chlorophyll a concentrations compared to shallow water bodies [1–4]. Likewise, lake size, depth and bathymetry determine the fraction of a lake’s habitat that can support photosynthetic carbon fixation and, thus, control the distribution of a lake’s growth limiting nutrients across living primary producer biomass and compartments at aphotic depths [5–7]. As a result, whole-lake primary production and its partitioning between pelagic and benthic producers are strongly dependent on lake morphometry and water transparency, with benthic production being largely determined by the fractional bottom area above the photosynthetic compensation point [5, 8–10]. Since the latter varies with the amount of shading from pelagic producers [11–15], predicting the productivity of lakes across large gradients in environmental conditions requires a process-based, mechanistic understanding of the dynamic interplay between light, nutrients, and photosynthetic biomass in relation to lake size, depth and basin shape.

The influence of physical drivers such as lake size, basin shape and water transparency on primary producers has been investigated theoretically using a variety of modelling approaches that can be categorized along three axes [16]: (1) Complexity of the included physical, chemical and biological ecosystem components, ranging from minimal models with one or two dynamic components [17, 18] to highly complex models with *>*100 state variables [19]; (2) Spatial dimensionality, ranging from simplified box-models [7, 20] to the integration of production modules into 3-dimensional hydrodynamical models [21, 22]; and (3) Degree to which dynamic feedback is taken into account, ranging from models that derive primary production from static empirical relationships with environmental factors [10, 23–25] to process-based, dynamical models that include feedbacks between primary producers and their environment [26–29] and hybrid models that use a combination of the two approaches [5, 22, 30].

While the rich body of modelling studies has yielded many insights into physical determinants of lake primary production, the sheer wealth of divergent modelling approaches has nevertheless hampered a more general and comprehensive understanding of the relative roles of these physical drivers across different types of lakes and their mechanistic interactions with relevant lake-internal processes. For example, a majority of more conceptual models of carbon and nutrient dynamics has focused exclusively on pelagic processes in highly simplified 0-or 1-dimensional vertical spatial settings [6, 31–33], thus neglecting the potential importance of the horizontal dimension, a varied bottom topography and benthic primary producers. On the other hand, models including benthic producers have often used static, empirical descriptions of certain processes rather than mechanistic representations [5, 10, 34], while models implemented in a fully 3-dimensional spatial setting have typically been tailored to specific systems [35–38], making it difficult to extract a more general, conceptual understanding of the role of lake morphometry in shaping whole-lake primary production. To our knowledge, a comprehensive theoretical investigation of the physical determinants of lake primary production from a conceptual, process-based perspective has not yet been performed.

In a first attempt to address this knowledge gap, we developed and analyzed ‘Lake2D’, a process-based, conceptual model that reduces lake bathymetry to its hypsographic depth distribution under the simplifying assumption of radial symmetry of the lake basin [39]. Lake2D adopts a 2-dimensional modeling approach that incorporates lake area, depth and hypsography, turbulent mixing in the horizontal and vertical directions, as well as benthic and pelagic algae and their interactions with dissolved nutrients and the sediment. In the present study, we use Lake2D to explore and analyze the dynamics of pelagic and benthic algae on a whole-lake scale in response to six environmental drivers (lake area and depth, horizontal and vertical turbulent mixing, water transparency and nutrient status), which we vary comprehensively over ranges that are representative of the vast majority of the world’s inland waters. We address the following specific questions: How do physical environmental conditions, including lake bathymetry, turbulent mixing, water transparency, and nutrient status, affect

- the total biomass of pelagic and benthic algae at the whole-lake scale?
- the relative contributions of benthic and pelagic algae to total biomass and the partitioning of a lake’s total nutrients between living biomass, dissolved and sedimented fractions?
- the spatial distribution of algal production and biomass, dissolved nutrients and detrital sediment across a lake’s hypsographic profile?

By answering these questions, we aim to identify general patterns and advance our conceptual understanding of the processes shaping primary production in lakes, providing theoretical insights and expectations about potential impacts of environmental change on different types of lakes.

## 2 Model description

In Section 2.1, we first present a 3-dimensional (3D) model of the dynamics of pelagic and benthic algae in a lake with arbitrary morphometry. From the 3D model, we derive in Section 2.2 a 2D simplification, Lake2D, which is the computational model used in this paper.

### 2.1 Full 3D model

The 3-dimensional model consists of the following five state variables: The concentrations of pelagic algal biomass *A* and a potentially growth-limiting inorganic nutrient *N*_d_ in the lake water, the areal densities of sedimented particular nutrient *N*_s_ at the lake bottom and of benthic algal biomass *B* at the sediment-water interface, and the intensity of the potentially growth limiting light *I* in the water column. Benthic and pelagic algal biomass is expressed in units of carbon, and the dissolved and sedimented nutrient is assumed to be phosphorus. The rates of change of these state variables are governed by four differential equations and one algebraic equation:

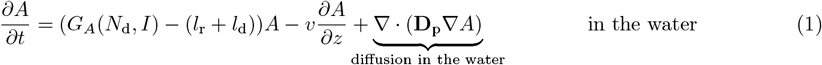

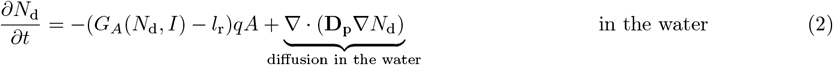

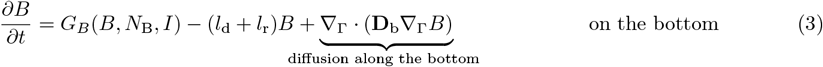

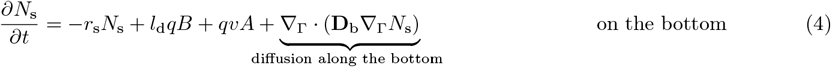

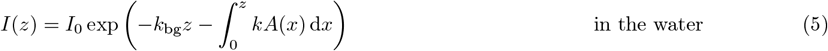

Here *z* is the depth below the lake surface and the diffusion terms are written in coordinate-free matrix-vector form which are labeled for clarity. The growth rates of pelagic and benthic algae, *G*_*A*_, and *G*_*B*_, are given by

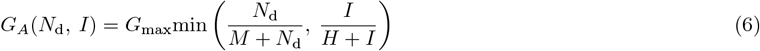

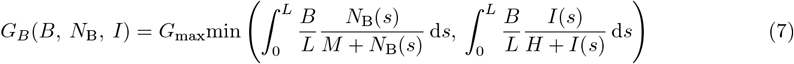

Equations 1–7 are explained in detail below. All variables and parameters are listed with their units in Table 1, using values based on previous work [6, 29, 32].

**Table 1.** State variables and parameters.

| Symbol | Unit | Value | Description |
| --- | --- | --- | --- |
| State variables |  |  |  |
| $A$ | $\text{mg C m}^{-3}$ | ... | Pelagic algal biomass concentration |
| $B$ | $\text{mg C m}^{-2}$ | ... | Benthic algal surface density |
| $I(z)$ | $\mu\text{mol photons m}^{-2}\text{s}^{-1}$ | ... | Light intensity in the lake water |
| $I(s)$ | $\mu\text{mol photons m}^{-2}\text{s}^{-1}$ | ... | Light intensity in the benthic algal layer |
| $N_B$ | $\text{mg P m}^{-3}$ | ... | Dissolved nutrient concentration in the benthic algal layer |
| $N_d$ | $\text{mg P m}^{-3}$ | ... | Dissolved nutrient concentration |
| $N_s$ | $\text{mg P m}^{-2}$ | ... | Benthic sedimented nutrient surface density |
| Variable environmental parameters |  |  |  |
| $d_r$ | $\text{m}^2 \text{ day}^{-1}$ | ... | Horizontal diffusion coefficient (2D-model) |
| $d_R$ | $\text{day}^{-1}$ | [0.001, 10] | Horizontal mixing rate |
| $d_z$ | $\text{m}^2 \text{ day}^{-1}$ | ... | Vertical diffusion coefficient |
| $d_Z$ | $\text{day}^{-1}$ | [0.001, 100] | Vertical mixing rate |
| $k_{bg}$ | $\text{m}^{-1}$ | 0.3, 1, 2 | Background light attenuation coefficient |
| $N_{\text{area}}$ | $\text{mg P m}^{-2}$ | 100, 300, 1000, 3000 | Total nutrient content per unit surface area |
| $r$ | m | ... | Distance from lake center in radial direction |
| $R$ | m | ... | Lake surface radius |
| $z$ | m | ... | Vertical distance from the water surface |
| $z_m$ | m | 1, 3, 10, 30 | Lake mean depth (2D-model), or max depth (1D-model) |
| Constant parameters and variables |  |  |  |
| $d_b$ | $\text{m}^2 \text{ day}^{-1}$ | $10^{-6}$ | Benthic orthogonal diffusion coefficient (2D-model) |
| $d_{N_B}$ | $\text{m}^2 \text{ day}^{-1}$ | $1.7 \times 10^{-4}$ | Benthic perpendicular diffusion coefficient |
| $f$ | $\text{mg C mg}^{-1}$ | 0.1 | Benthic carbon mass to wet weight ratio |
| $G_{\text{max}}$ | $\text{day}^{-1}$ | 1.5 | Maximum specific growth rate |
| $I_0$ | $\mu\text{mol photons m}^{-2}\text{s}^{-1}$ | 300 | Light intensity at lake surface |
| $k$ | $\text{m}^2\text{mg C}^{-1}$ | $3 \times 10^{-4}$ | Light attenuation coefficient of algal biomass |
| $L$ | m | ... | Thickness of the benthic algal layer |
| $M$ | $\text{mg P m}^{-3}$ | 4 | Half saturation constant of nutrient-dependent algal production |
| $H$ | $\mu\text{mol photons m}^{-2}\text{s}^{-1}$ | 60 | Half-saturation constant of light-dependent algal production |
| $l_d$ | $\text{day}^{-1}$ | 0.02 | Mortality loss rate of algae |
| $l_r$ | $\text{day}^{-1}$ | 0.08 | Respiration loss rate of algae |
| $q$ | $\text{mg P mg C}^{-1}$ | 0.015 | Algal stoichiometric P:C ratio |
| $\rho_w$ | $\text{mg m}^{-3}$ | $997 \times 10^6$ | Density of water |
| $r_s$ | $\text{day}^{-1}$ | 0.02 | Mineralization rate of sedimented nutrients |
| $v$ | $\text{m day}^{-1}$ | 0.1 | Algal sinking speed |
| $\Gamma$ | — | - | Lake bottom surface |
| $\Omega$ | — | - | Lake volume |

**Table 2.**
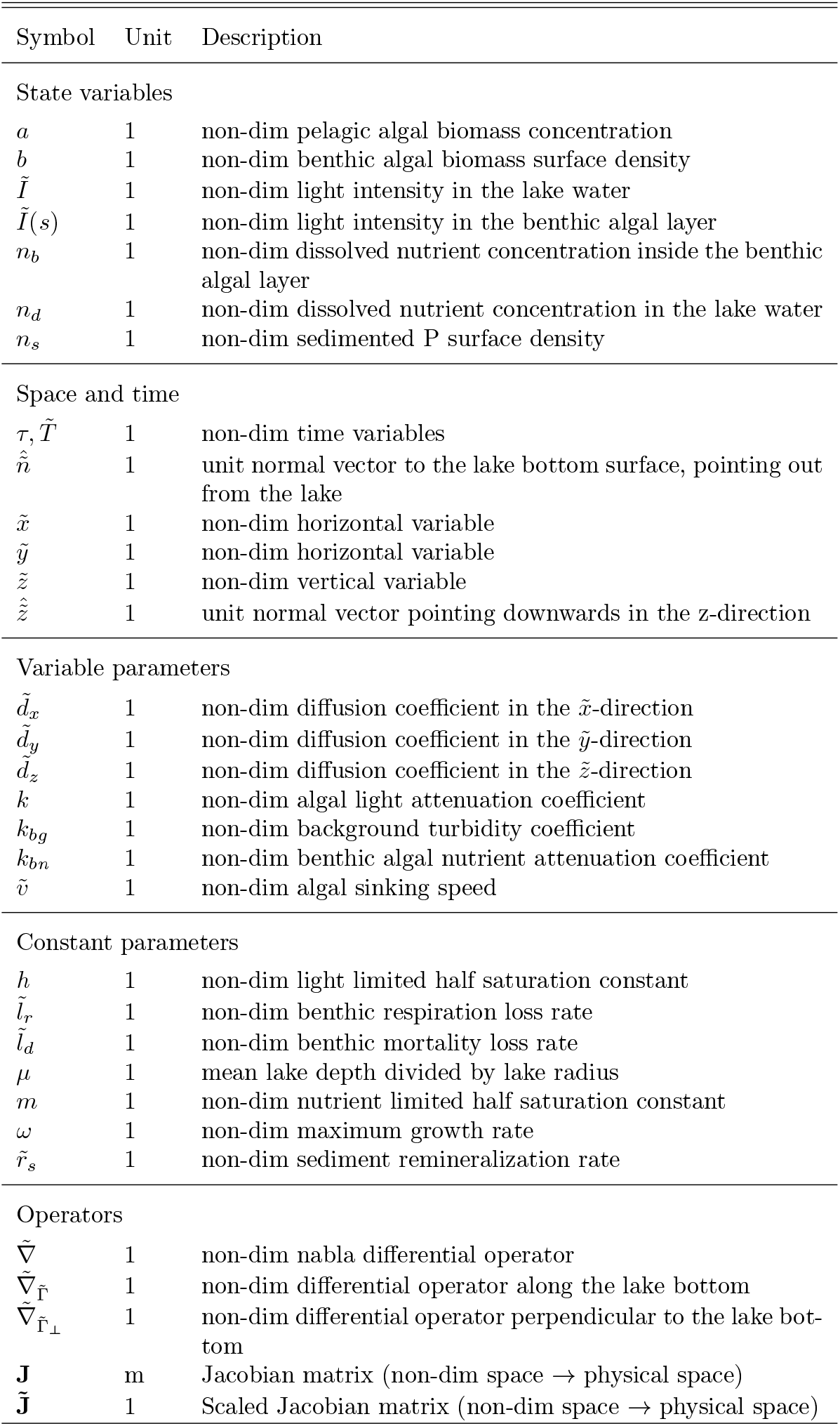
Non-dimensionalized state variables, parameters, operators, and vectors.

#### Transport mechanisms in the water

The model considers two spatial transport processes in the water column, turbulent mixing and sinking. Turbulent mixing of pelagic algae and dissolved nutrients is described by vertical and horizontal diffusion. In addition, pelagic algae sink at a constant velocity *v*. Pelagic algae that reach the bottom die and their nutrients turn into sedimented particular nutrients. Benthic algae and sedimented nutrients experience very slow diffusion along the bottom. We furthermore assume that the system is closed for nutrients, leading to the zero-flux boundary conditions

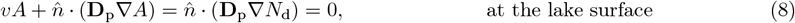

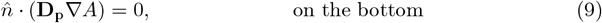

where 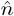 is the outward-pointing normal vector to each surface. The terms 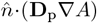 and 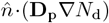 are the diffusive fluxes (mg m^−2^ day^−1^) of *A* and *N*_d_ across the denoted surfaces. The boundary condition for dissolved nutrients at the bottom depends on several cross-boundary processes described further down.

#### Pelagic algal dynamics

The growth of pelagic algae at a given position in the lake is either limited by the local light intensity, or the local dissolved nutrient concentration, as described by the minimum function of two Monod terms in (6) where *G*_max_ is the maximum specific growth rate and *H* and *M* are the half-saturation constants of light and nutrient-limited growth, respectively. Light intensity *I* attenuates vertically in the water column according to (5), where pelagic algae contribute with specific attenuation coefficient *k* and background attenuation with coefficient *k*_bg_. The latter represents light attenuation by non-algal materials in the water and by water itself. In addition to sedimentation losses, pelagic biomass is lost through background respiration and mortality with fixed rates *l*_r_ and *l*_d_, respectively.

#### Pelagic nutrient dynamics

Uptake and recycling of dissolved nutrients at a given depth are driven by local algal growth and losses. We assume that the stoichiometric phosphorus-to-carbon (P:C) ratio *q* of algal biomass is fixed. Consequently, the local uptake of dissolved phosphorus equals a fraction *q* of algal carbon production. Similarly, algal (carbon) biomass losses through mortality and respiration are accompanied by proportional phosphorus losses, which we assume to be instantly recycled in dissolved form.

#### Sediment nutrient dynamics

Particular nutrients enter the sediment through two processes – sinking of pelagic algae and background mortality of benthic algae at the rate *l*_d_. We assume that remineralization of sedimented nutrients is a first order process at the rate *r*_s_ and that remineralized nutrients are released into the water directly above the bottom (see (14) below).

#### Benthic algal dynamics

To focus our analyses on the impact of physical factors, we assume that benthic and pelagic algae have identical traits for growth and metabolism. Thus, benthic and pelagic algae have identical maximum specific growth rates and half saturation constants for light- and nutrient-dependent growth, identical light attenuation coefficients and elemental P:C ratios, as well as identical respiration and mortality rates.

To calculate their growth rate, we assume that benthic algae form a homogeneous layer of thickness *L*, which depends on benthic algal biomass as

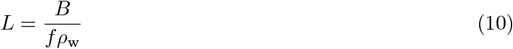

where *f* is the ratio of benthic algal carbon to wet mass and *ρ*_w_ is the specific density of algal wet mass assumed to be identical to the density of water. Light in the benthic algal layer is attenuated according to Lambert-Beer’s law. Assuming that non-algal light attenuation is negligible, light-limited benthic algal growth can then be integrated over the benthic algal layer as [40]

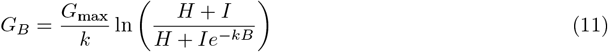

where *I* is the local light intensity at the bottom of the water column according to (5).

Nutrient-limited benthic algal growth is modeled analogously by integrating over the vertical profile of dissolved nutrients in the benthic algal layer. We assume that dissolved nutrients immediately above the lake bottom diffuse into the benthic algal layer while simultaneously being consumed by benthic algae. These two processes are modeled with the differential equation

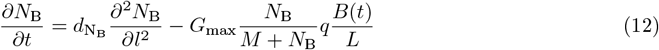

where *N*_B_ is the concentration of dissolved nutrient in the benthic algal layer, 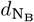 is the diffusion co-efficient in the benthic algal layer, *l* ∈ [0, *L*] denotes the orthogonal position inside the benthic algal layer, and *q* is the P:C ratio of benthic biomass. We assume that these processes are fast compared to other processes, and therefore treat (12) on a separate timescale. For each time *t*, we approximate the steady state solution of (12) by linearizing the benthic algal consumption term and solving the resulting equation analytically. We further assume that the nutrient concentration at the surface of the benthic algal layer equals the concentration at the bottom of the water column and that nutrients do not diffuse out of the bottom of the benthic algal layer, yielding the boundary conditions

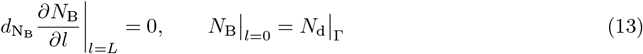

where Γ is the lake bottom and 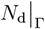 is the dissolved nutrient concentration along the bottom of the lake. We then integrate the nutrient-limited growth over the benthic algal layer (first integral in (7)), using the approximated solution of (12) for each time step. In a final step, benthic algal growth per unit area is calculated as the minimum of nutrient and light-dependent growth (7). This approach can sometimes overestimate benthic algal growth, since it is possible that growth switches from nutrient to light limitation somewhere inside the benthic algal layer. Note that *N*_B_ is a dummy variable that is only used to calculate nutrient-limited benthic algal growth and therefore does not contribute to the nutrient mass balance of the system.

#### Nutrient fluxes at the sediment-water interface

We assume that nutrients lost through benthic algal respiration are released in dissolved form into the water directly above, which is the same water from which the nutrients are drawn that sustain benthic algal growth. Together with the release of remineralized particular nutrients from the sediment, this yields the following boundary condition for dissolved nutrients at the bottom of the water column

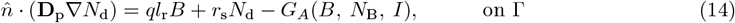

#### Nutrient conservation

The total nutrient content of a modeled lake is defined as total phosphorus per unit surface area

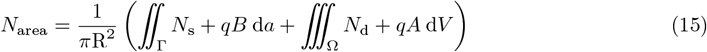

where Γ is the bottom surface, and Ω is the lake volume.The absence of internal nutrient sources and the closed boundary conditions imply that *N*_area_ is a measure of nutrient enrichment, which is determined by the initial conditions and remains constant over time.

### 2.2 Model Lake2D

Lake2D is derived from the full 3-dimensional model described in Section 2.1 using the following simplifying assumptions on lake geometry and symmetries on the problem. We choose a cone-shaped, radially symmetric lake geometry because its simple topography facilitates visualization and interpretation of results, and also because its bottom profile is similar to the hypsography of many real lakes [41–43]. We assume that the intensities of horizontal and vertical mixing are independent of each other and do not vary in space or time. In addition, we also assume radial symmetry such that for any distance from the center vertical axis of the lake, all state variables are well-mixed in the tangential direction. This implies that transport processes described by horizontal diffusion operate exclusively in the radial direction, and that spatial dynamics can be fully described with a 2-dimensional implementation of horizontal and vertical transport processes.

#### Model implementation

To facilitate execution and analysis, we non-dimensionalized the model (Appendix A) and implemented it on a half cross-section of the lake. The full 3D cone-shaped geometry can be retrieved by rotating this triangular cross-section around the center axis of the lake. The radial symmetry of this approach implies a reflective boundary condition for pelagic algae and dissolved nutrients along the left boundary of the triangular cross-section, corresponding to the center of the lake:

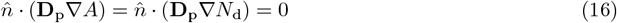

where 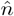 is the unit normal vector to the vertical center line of the lake, pointing outwards in the horizontal plane. In cylindrical coordinates, this boundary condition can be written as

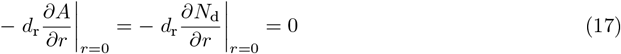

where *r* is the distance from a point to the center of the lake in the horizontal plane.

## 3 Methods

### Design of numerical experiments

We conducted extensive numerical simulations of Lake2D to evaluate algal performance across a wide range of environmental conditions. Six key parameters were systematically varied: background attenuation coefficient (*k*_bg_), mean depth (*z*_m_), total nutrient content per surface area (*N*_area_), lake radius (*R*), horizontal diffusion coefficient (*d*_*r*_), and vertical diffusion coefficient (*d*_*z*_). To reduce the dimensionality of the parameter space, we combined the horizontal diffusion coefficient and lake radius into a single parameter, the horizontal mixing rate *d*_R_ defined as

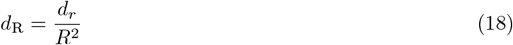

This parameter effectively captures the essential physics of horizontal mixing and lake size while simplifying the overall analysis. We analogously combined the mean depth and the vertical diffusion coefficient into the vertical mixing rate *d*_Z_

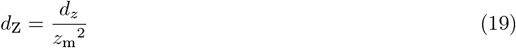

Together, *d*_R_ and *d*_Z_ constitute a two-dimensional slice of the parameter space that we hereafter refer to as the ‘mixing space’.

Note that the effects of varying *R* and *d*_r_ are fully captured by the horizontal mixing rate *d*_R_, such that a given change in the value of *d*_R_ can be interpreted as a change in either lake surface area or the intensity of horizontal mixing. Because the horizontal dimension of most real lakes exceeds their vertical dimension by several orders of magnitude, while the intensity of turbulent mixing is typically several orders of magnitudes higher in horizontal than in vertical direction [44], we interpret the horizontal mixing rate *d*_R_ as an inverse measure of lake area. Similarly, a given change in the vertical mixing rate *d*_Z_ can be interpreted as a change in either lake mean depth or the intensity of vertical mixing. However, since pelagic algae sink at a constant velocity, *d*_Z_ does not capture all effects of lake depth, and *z*_m_ needs to be treated as an independent parameter.

Collectively, *z*_m_, *k*_bg_, *N*_area_, and the mixing space define a 5-dimensional parameter space, which we numerically explored in a fully factorial design, where the parameters *k*_bg_, *z*_m_, and *N*_area_ were varied over ranges that are representative of the vast majority of the world’s lakes, ponds, and reservoirs [10, 45–47]. Specifically, we performed 19200 numerical simulations resulting from the combination of four values of *N*_area_ (100, 300, 1000, and 3000 mgP m^−2^), four values of *z*_m_ (1, 3, 10, and 30 m), two values of *k*_bg_ (0.3, 1 m^−1^), and a 20 × 20 factorial mixing space, where the horizontal and vertical mixing rates *d*_R_ and *d*_Z_ were varied on a logarithmic scale over the intervals [0.001, 10] day^−1^ and [0.001, 100] day^−1^, respectively.

### Execution of numerical experiments

We implemented the non-dimensionalized version of Lake2D in MATLAB using a finite volume method, and the built-in function ODE15s was used to integrate in time. A square grid with a minimum resolution of 30×30 equally spaced grid cells was employed, and the grid was cut diagonally resulting in triangular grid elements along the bottom of the lake.

Under most environmental conditions, steep spatial gradients could arise at specific depths or distances from the lake center. In such cases, the local grid resolution near the steep gradients was increased as necessary to maintain accuracy. All simulations were run to steady state from the same initial condition: 98 % of total nutrients in dissolved form, 1% in pelagic algae, and the remaining 1% in benthic algae, all evenly distributed across the lake’s volume and bottom, respectively. Across the explored 5-dimensional parameter space, all simulations converged to equilibrium steady state within 70-130 days. The only exceptions were very shallow (*z*_m_ = 1 m) nutrient-poor (*N*_area_ ≤ 300 mgP m^−2^) lakes, which took an order of magnitude longer to converge.

### Model output and visualization

We focus our analyses on model output at the whole-lake scale. For each individual model run, we therefore integrated steady-state pelagic, benthic, and total (pelagic + benthic) algal biomass over the entire lake and expressed it as average biomass per unit of lake surface area. To visualize the results in the 5-dimensional parameter space we proceeded as follows. For each combination of *z*_m_, *k*_bg_, and *N*_area_ values, we first produced heat maps of the steady-state pelagic, benthic, and total algal biomasses in *d*_R_ - *d*_Z_ mixing space (see Fig. 1 for representative examples).

**Figure 1:**
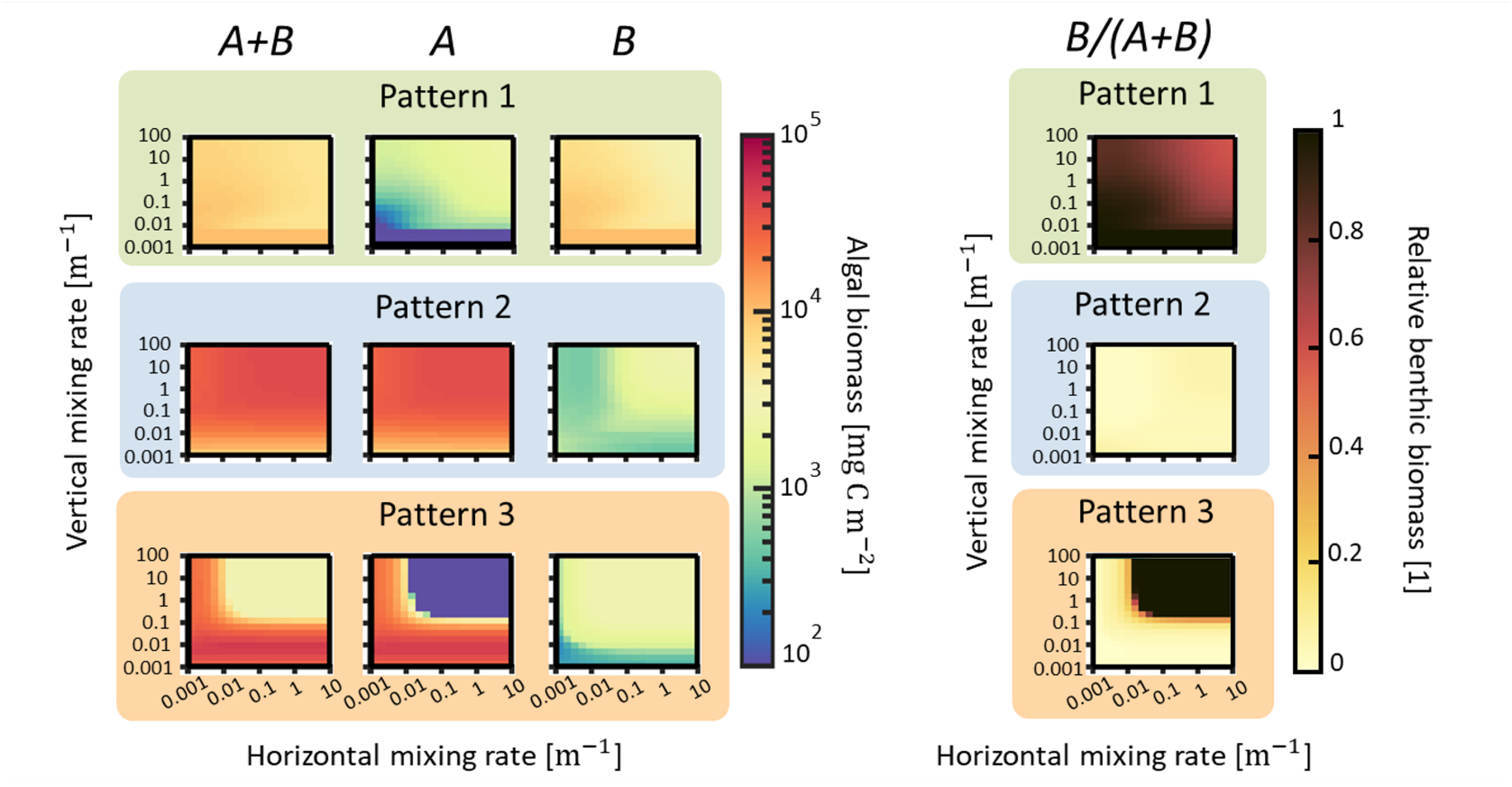
Heatmaps of lake-wide pelagic (A), benthic (B), total (A+ B), and relative benthic (B/(A+B)) algal biomass for patterns 1,2, and 3 in Fig. 2, marked with a green, blue, and orange background, respectively. Background attenuation k_bg_ = 1 m^−1^ in all panels. The remaining parameter values are N_tot_ = 300 mg P m^−1^, z_m_ = 1 m for pattern 1, N_tot_ = 1000 mg P m^−1^, z_m_ = 10 m for pattern 2, and N_tot_ = 3000 mg P m^−1^, z_m_ = 30 m for pattern 3. The horizontal mixing rate d_R_ varies along the horizontal axis of each panel in the range [0.001, 10] day^−1^, and the vertical mixing rate d_Z_ varies along the vertical axis of each panel in the range [0.001, 100] day^−1^. Both axes are plotted on a log10 scale. Each pixel in the panels of theA+B, A, and B columns is the average biomass per unit of lake surface area (mg C m^−2^) on a log10 scale. The relative benthic algal biomass (rightmost column of panels) is expressed on a linear scale.

Inspection of the heat maps revealed the existence of three qualitatively distinct patterns by which different combinations of the environmental parameters can affect spatially integrated pelagic, benthic, and total algal biomass, as well as the contribution of benthic to total algal biomass, at the whole-lake scale (Fig. 1–3). A deeper understanding of these patterns requires a closer examination of the within-lake spatial distributions of nutrients and algae under different mixing scenarios. To facilitate this, we selected one representative example of each of the three patterns for which we show individual simulation output from select areas in mixing space: a high mixing scenario, a high vertical - low horizontal mixing scenario, a low mixing scenario, and an intermediate mixing scenario.

## 4 Results

Depth-integrated biomass exhibits several trends across the 5-dimensional environmental parameter space (Fig. 2). To increase readability, conditions of low (*k*_bg_ = 0.3 m^−1^) and high (*k*_bg_ = 2 m^−1^) background turbidity were omitted from Fig. 2 since they are qualitatively largely redundant with *k*_bg_ = 1 m^−1^ (see Appendix B1 for a full figure). First, we describe and explain general trends in the responses of pelagic, benthic, and total algal biomass, as well as the contribution of benthic to total algal biomass, at the whole-lake scale to the environmental drivers nutrient content, background attenuation, lake depth, and mixing space. We subsequently explore and illustrate the underlying mechanisms further through inspection of within-lake spatial patterns in select simulations that cover a relevant range of environmental and mixing conditions.

**Figure 2:**
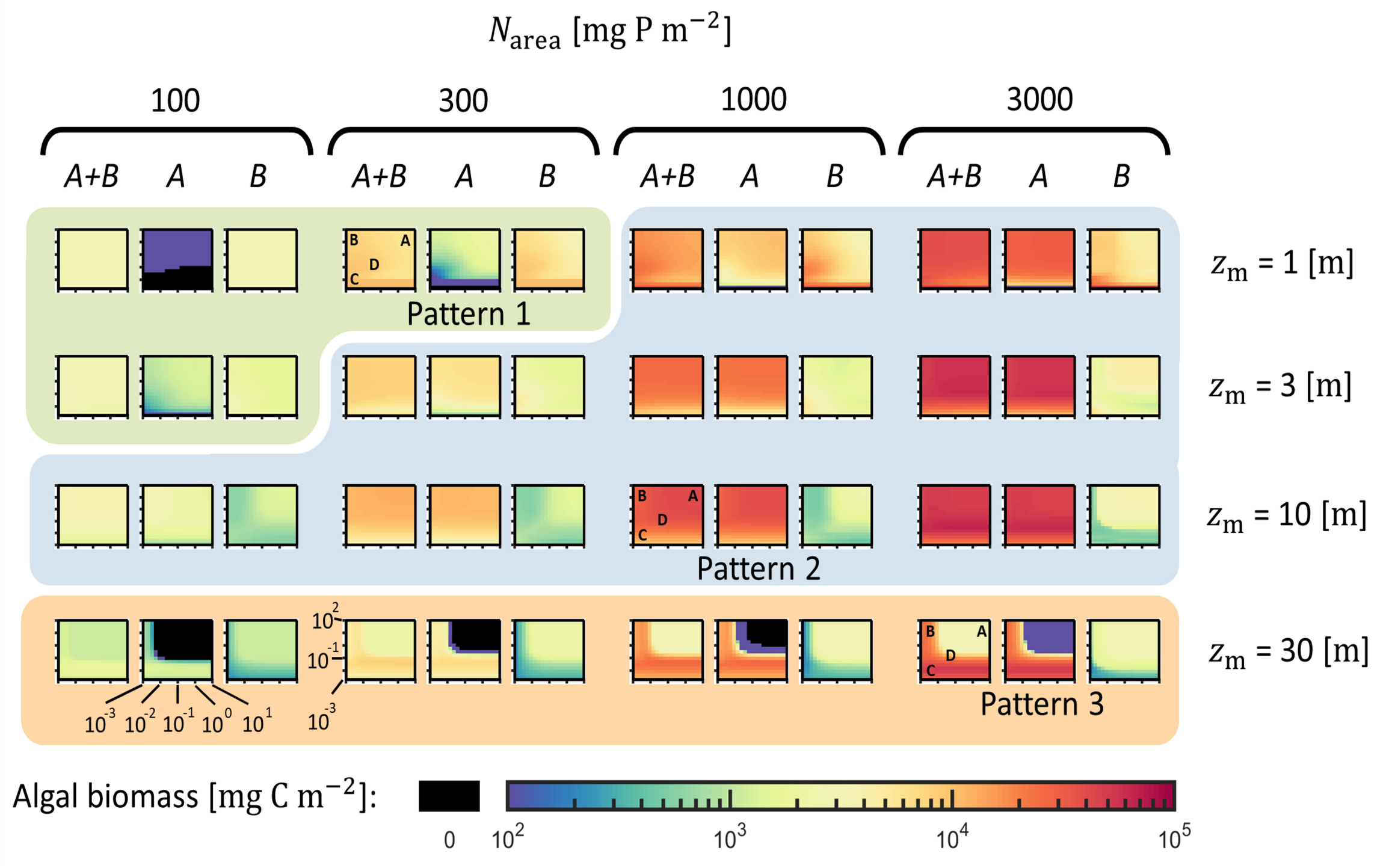
Heatmaps of lake-wide pelagic (A), benthic (B), and total (A+B) algal biomass for varying total nutrient content per surface area N_tot_, mean depth z_m_, and diffusive mixing rates d_R_ and d_Z_. Background turbidity k_bg_ = 1 m^−1^ in all panels. The horizontal mixing rate d_R_ varies along the horizontal axis of each panel in the range [0.001, 10] day^−1^, and the vertical mixing rate d_Z_ varies along the vertical axis of each panel in the range [0.001, 100] day^−1^. Both axes are plotted on a log10 scale. Each pixel in a panel is the average biomass per unit of lake surface area (mg C m^−2^) on a log10 scale (see color bar at the bottom of the figure). Black areas indicate extinction, defined as an average concentration < 0.001 mg C m^−3^. Panels on a green background show pattern 1, panels on a blue background show pattern 2, and panels on an orange background show pattern 3. Letters A-D inside panels highlight the mixing rate combinations of the simulations shown in Fig. 7–9.

### 4.1 Contributions of pelagic and benthic algae to total biomass

Pelagic algae make up the bulk of total biomass in most of the parameter space (see pattern 2-panels on blue background in Fig. 2, compare panels *A* with panels *A*+*B*), with three exceptions: (1) at extremely low vertical mixing rates (largely outside the range shown in Fig. 2) where sinking losses drive pelagic algae extinct; (2) in shallow, nutrient-poor lakes where nutrient-limited pelagic algal growth cannot compensate for high sinking losses (pattern 1-panels on green background); and (3) in deep lakes where high mixing to aphotic depths causes severe light limitation of pelagic algae (pattern 3-panels on orange background). In the latter two cases, benthic algae dominate total biomass in either all or parts of the mixing space (dark brown to black areas in Fig. 3). Environmental conditions where pelagic and benthic algae contribute more equally to total biomass are almost absent (*N*_area_ = 1000, *z*_m_ = 1 in Fig. 2, 3). Total biomass therefore shows near identical trends to benthic biomass in most pattern 1 lakes and in pattern 3 lakes at high overall mixing, and near identical trends to pelagic biomass in most pattern 2 lakes and in pattern 3 lakes at low horizontal and/or vertical mixing.

**Figure 3:**
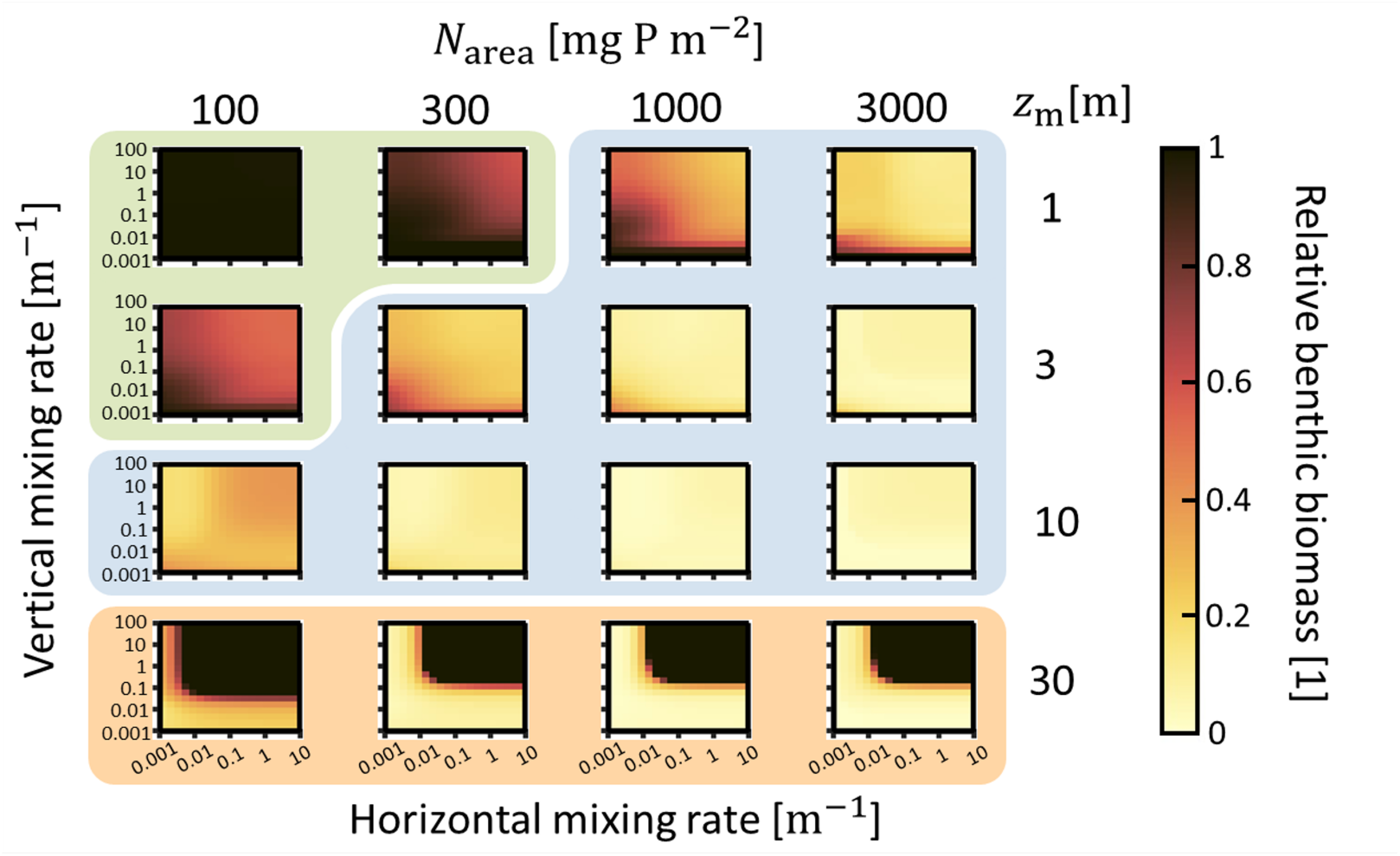
Heat maps of lake-wide benthic biomass divided by total biomass for varying background turbidity k_bg_, total nutrient content per surface area N_tot_, mean depth z_m_, and diffusive mixing rates d_R_ and d_Z_. The horizontal diffusion mixing rate d_R_ is varied along the horizontal axis of each panel in the range [0.001, 10] day^−1^, and the vertical diffusion mixing rate d_Z_ is varied along the vertical axis of each panel in the range [0.001, 100 day^−1^. Both axes are plotted on a log10 scale. Each pixel in a panel is the unitless ratio of the average benthic biomass divided by the total biomass, see the colorbar on the right side of the figure. Panels with a green background are pattern 1 sweeps, panels with a blue background are pattern 2 sweeps, and panels with an orange background are pattern 3 sweeps.

### 4.2 Varying nutrient content per unit area *N*_area_

Of the environmental parameters explored, total nutrient content per area (*N*_area_) has the largest impact on both pelagic and total algal biomass, while having only a weak effect on benthic biomass in all but the shallowest and most nutrient poor systems (Fig. 2). Because of the logarithmic biomass scale of Fig. 2, the impact of nutrient enrichment becomes more visible in a difference plot (Fig. 4), which illustrates a systematic difference in resource limitation between pelagic and benthic algae. Specifically, algal responses to nutrient enrichment suggest that lake-integrated pelagic algal biomass is primarily nutrient-limited (blue areas), while benthic algal biomass is primarily light-limited (white areas). The latter becomes obvious in shallow, nutrient-rich systems, where nutrient enrichment even has a negative effect on benthic biomass because of strong shading from increased pelagic biomass (upper right panels in Fig. 4). While the results in Fig. 4 are for an intermediate background attenuation of *k*_bg_ = 1 m^−1^, very similar patterns are observed for *k*_bg_ = 0.3 and 2 m^−1^ (not shown).

**Figure 4:**
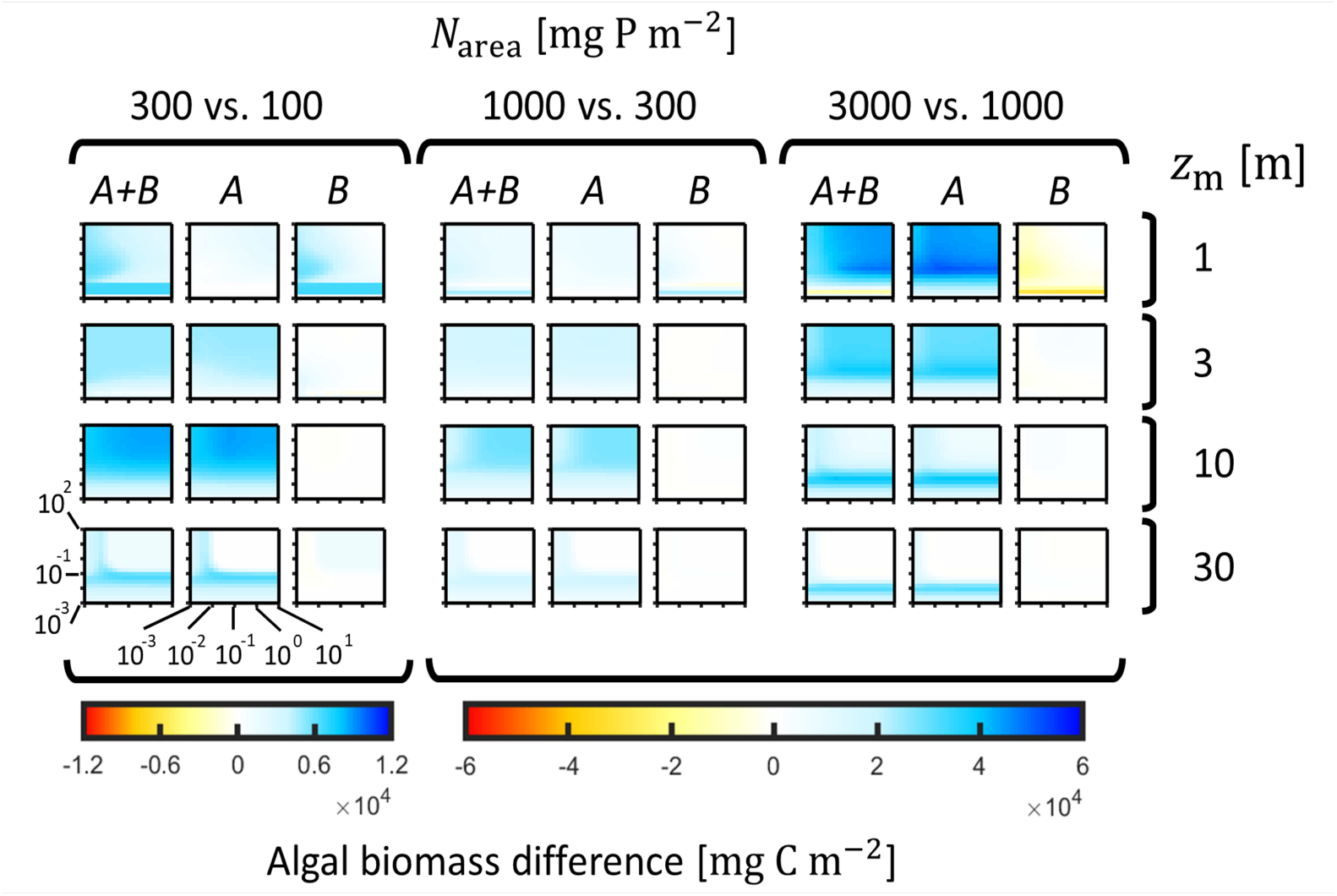
Differential heatmaps illustrating the impact of increasing nutrient content (N_area_) on algal biomass. Each panel displays the pixel-wise difference between two heatmaps from Fig 2, where a heatmap with a lower Narea value is subtracted from the heatmap with the next-higher Narea value, with all other parameters equal. For example, the A, B, and A+B panels in the top left corner visualize changes in algal biomass when Narea is increased from 100 to 300 mg P m^−2^ for z_m_ = 1 m. k_bg_ = 1 m^−1^ in all panels.

### 4.3 Varying background attenuation *k*_bg_

With increasing background attenuation (*k*_bg_), light availability decreases monotonically, which should have a negative effect on biomass when algal growth is primarily light-limited. In a difference plot comparing intermediate with low *k*_bg_, such a pattern is seen in deep lakes (*z*_m_ = 30 m) under conditions of high mixing to aphotic zones, where pelagic algae decline sharply with an increase in *k*_bg_ (bottom row of panels in Fig. 5). Under these same conditions, benthic algae show a slight compensatory increase in biomass caused by reduced shading from pelagic algae.

**Figure 5:**
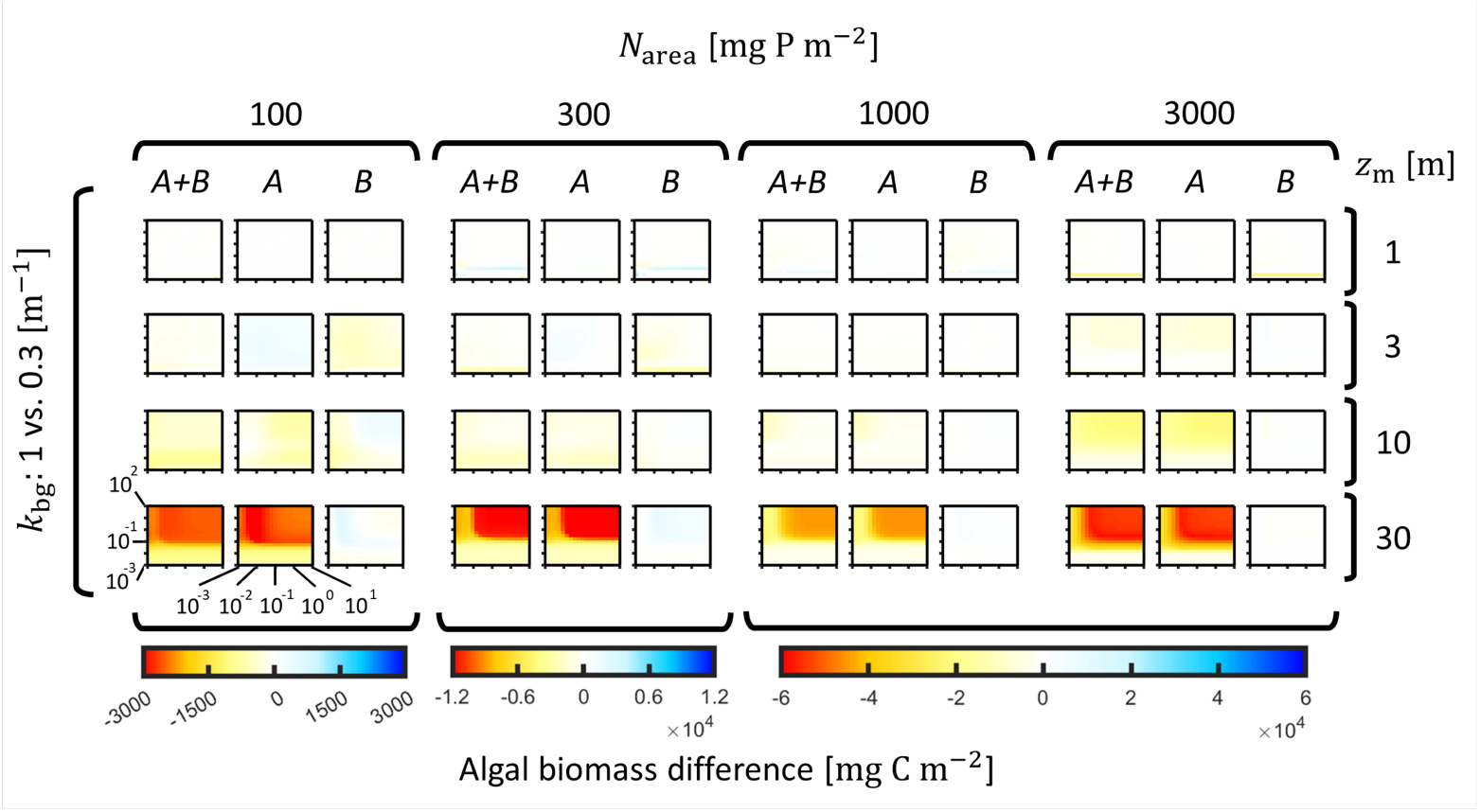
Differential heatmaps illustrating the impact of increasing background attenuation (k_bg_) on algal biomass. Each panel displays the pixel-wise difference resulting from the subtraction of a heatmap with a lower k_bg_ value from the heatmap with the next-higher k_bg_ value, with all other parameters equal. For example, the A, B, and A+B panels in the top left corner visualize changes in algal biomass when k_bg_ is increased from 0.3 to 1 m − 1 for N_area_ = 100 mg P m^−2^ and z_m_ = 1 m.

Yet, over most of the remaining parameter range, the negative effect of *k*_bg_ on lake-integrated algal biomass is relatively small, because pelagic algae tends to be primarily nutrient limited and benthic algae persist at shallower depths when light becomes more limiting. At higher levels of *k*_bg_ ≥ 2 m^−1^, the negative effect of background attenuation on pelagic biomass under high mixing becomes, however, increasingly relevant also in shallower lakes since light attenuates as the product *k*_bg_ · m (Fig. Appendix B1).

### 4.4 Varying mean depth *z*_m_

Of the environmental parameters explored, lake mean depth (*z*_m_) has the most complex impact on algal biomass (Fig. 6). This complexity arises from the interplay of three phenomena that have opposite effects on biomass. All else being equal, increasing *z*_m_ reduces (1) pelagic algal sinking losses, since algae must travel a longer distance before reaching the lake bottom; (2) average light intensity in the water and at the lake bottom, since a larger fraction of light is lost to background attenuation; (3) average nutrient concentration, since a given amount of nutrient per area (*N*_area_) is diluted in a larger volume of water. Consequently, pelagic biomass peaks at intermediate lake depths and declines towards both shallower lakes (where sinking losses are high) and deeper lakes (where light is strongly limiting) (Fig. 2, 6). In contrast, since benthic algae do not experience sinking losses, benthic biomass declines uniformly with lake depth, with the exception of the deepest, well-mixed lakes where benthic algae benefit from the extinction of pelagic algae (Fig. 2, 6). Again, the results in Fig. 6 are for a background attenuation of *k*_bg_ = 1 m^−1^, but similar patterns are observed for *k*_bg_ = 0.3 and 2 m^−1^ (not shown).

**Figure 6:**
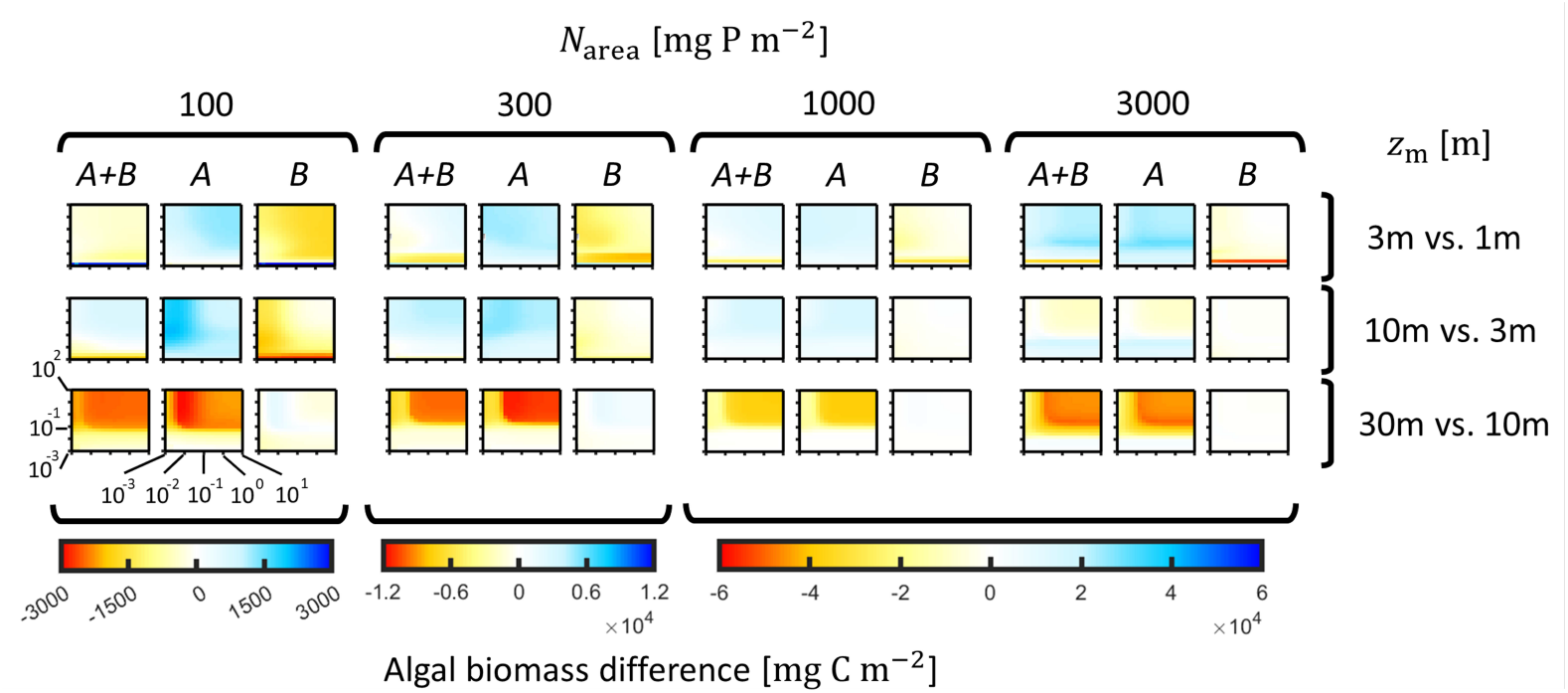
Differential heatmaps illustrating the impact of increasing mean depth (z_m_) on algal biomass. Each panel displays the pixel-wise difference between two heatmaps from Fig. 2 where a heatmap with a lower z_m_ value is subtracted from the heatmap with a higher z_m_ value, with all other parameters equal. For example, the A, B, and A+B panels in the top left corner visualize changes in algal biomass when z_m_ is increased from 1 to 3 m, N_area_ = 100 mg P m^−2^ and k_bg_ = 0.1 m^−1^. k_bg_ = 1 m^−1^ in all panels.

### 4.5 Varying horizontal and vertical mixing *d*_R_ **and** *d*_Z_

The influence of horizontal and vertical mixing on algal biomass varies systematically along the above-described gradients in nutrient content, background attenuation and lake mean depth. We illustrate these trends by focusing on three representative lake types, i.e. one example each of a shallow, nutrient-poor lake (pattern 1), a deeper and/or nutrient-rich lake (pattern 2), and a very deep and/or highly turbid lake (pattern 3) (Fig. 1, 2). Below we first give an overview of the partitioning of the system’s nutrients among the four state variables (*A, B, N*_d_, *N*_s_) in pattern 1-3 lakes. Starting from a well-mixed reference scenario, we then describe the influences of reduced horizontal and vertical mixing on the four state variables and their within-lake distribution in the three lake types. We end by describing why pelagic and benthic algae show diagonal trends in mixing space that are anti-correlated in pattern 1 and 3 lakes, but correlated in pattern 2 lakes.

#### 4.5.1 Nutrient partitioning in pattern 1-3 lakes

In shallow, nutrient-poor pattern 1 lakes, pelagic algae attain relatively low growth rates and experience very high sinking losses. Pelagic biomass is therefore considerably lower than benthic biomass under all mixing conditions, dissolved nutrient concentrations are very low, algal growth is nutrient-limited, and the majority of the system’s nutrients is stored in sediment and benthic algae (Fig. 7A-D). In contrast, in more nutrient-rich and/or deeper pattern 2 lakes, pelagic algal growth and sinking loss rates are more balanced. Pelagic biomass is therefore considerably higher than benthic biomass under all mixing conditions (Fig. 8A-D). Most production occurs at shallow depths where nutrients are limiting, while light is strongly limiting at greater, aphotic depths where a sizable fraction of the system’s nutrients is present in dissolved form. Finally, in very deep and/or highly turbid pattern 3 lakes, light limitation is further exacerbated to the degree that pelagic algae go extinct when vertical and horizontal mixing is high. This creates complementary L-shapes of pelagic and benthic biomass in mixing space (bottom row of panels in Fig. 2). Due to strong light limitation, total algal biomass and sediment nutrients are, however, low and the majority of the system’s nutrients is present in dissolved form (Fig. 9A-D).

**Figure 7:**
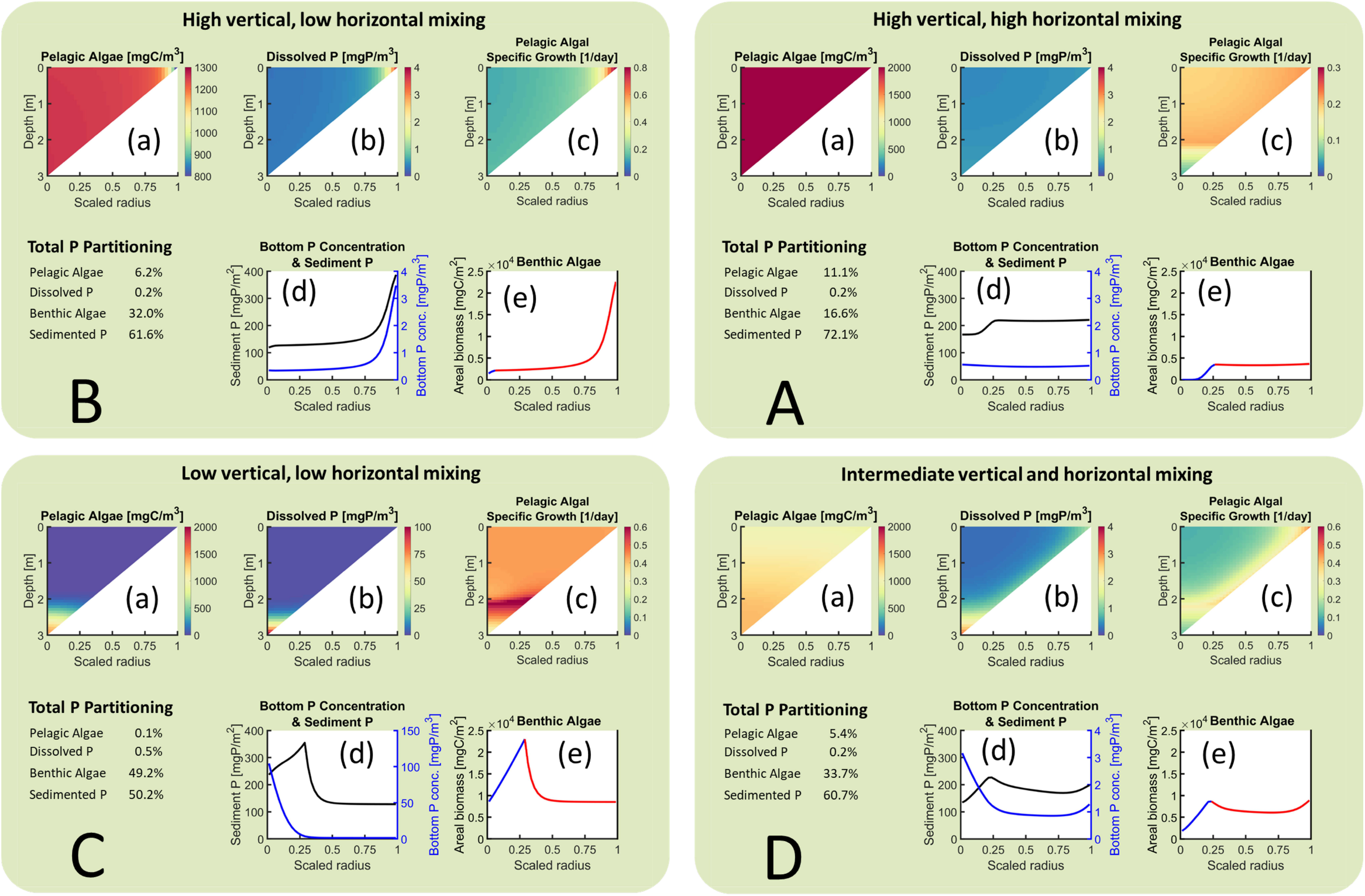
(Previous page.) Four example simulations visualizing 2D model output that yields pattern 1 in Fig. 2. Each panel A-D displays one 2D simulation selected from the top left, top right, bottom left, and middle of mixing space, see the markings A-D in Fig. 1 (panels with green background). In each panel, the top row of triangular heatmaps displays the concentration of pelagic algae (a), concentration of dissolved phosphorus (b), and pelagic algal specific growth rate (c) respectively. The vertical left edge in sub-panels (a)-(c) corresponds to the center of the lake, and the x-axis is the relative distance from the lake center. Sub-panel (d) in the bottom center displays the sedimented phosphorus surface density (black) and the dissolved phosphorus concentration along the bottom (blue) Sub-panel (e) in the bottom right displays the surface density of benthic alge where the color of the line indicates if the benthic algal growth is light-limited (blue) or nutrient-limited (red). Note that the scales of the y-axes and color bars can differ between panels A-D. The table of values in the bottom left of each panel A-D lists the percentage of total phosphorus in each state variable.

**Figure 8:**
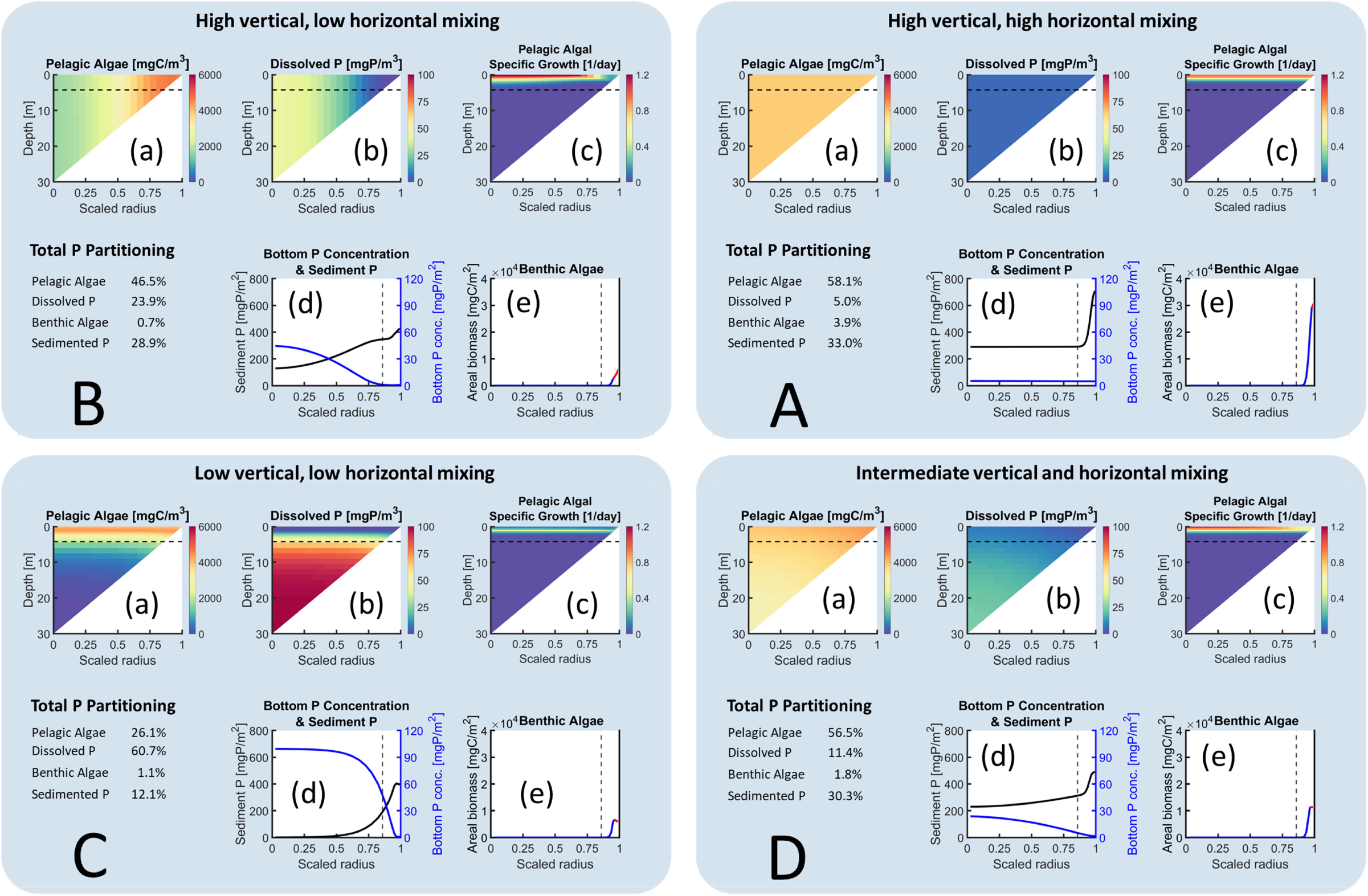
(Previous page.) Four example simulations visualizing 2D model output that yields pattern 2 in Fig. 2. Each panel A-D displays one 2D simulation selected from the top left, top right, bottom left, and middle of mixing space, see the markings A-D in Fig. 1 (panels with green background). In each panel, the top row of triangular heatmaps displays the concentration of pelagic algae (a), concentration of dissolved phosphorus (b), and pelagic algal specific growth rate (c) respectively. The vertical left edge in sub-panels (a)-(c) corresponds to the center of the lake, and the x-axis is the relative distance from the lake center. The compensation depth is marked in (a)-(c) with a horizontal dotted line. Sub-panel (d) in the bottom center displays the sedimented phosphorus surface density (black) and the dissolved phosphorus concentration along the bottom (blue) Sub-panel (e) in the bottom right displays the surface density of benthic alge where the color of the line indicates if the benthic algal growth is light-limited (blue) or nutrient-limited (red). The compensation depth is marked in (d)-(e) with a vertical dotted line. Note that the scales of the y-axes and color bars can differ between panels A-D. The table of values in the bottom left of each panel A-D lists the percentage of total phosphorus in each state variable.

**Figure 9:**
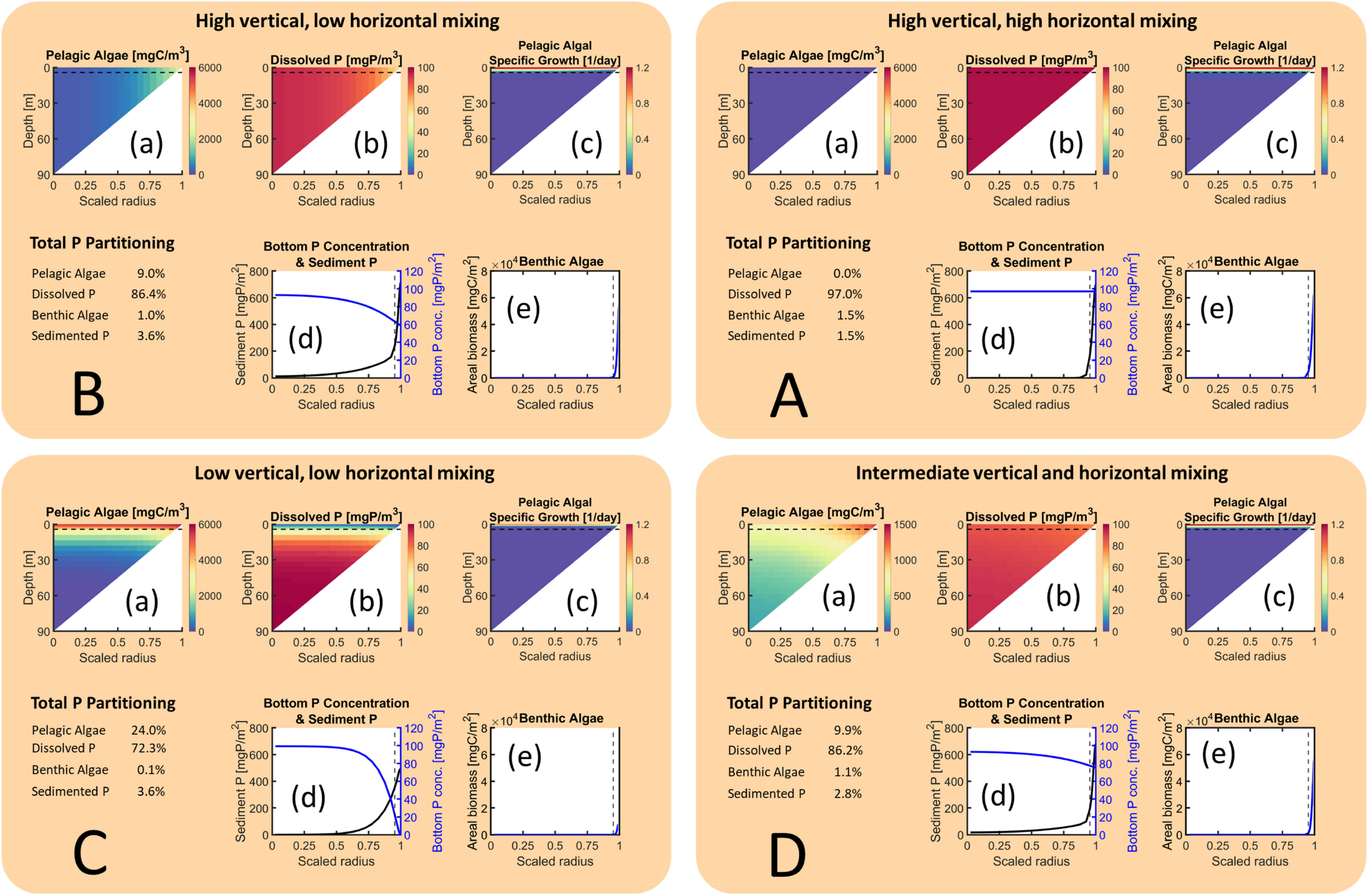
(Previous page.) Four example simulations visualizing 2D model output that yields pattern 3 in Fig. 2. Each panel A-D displays one 2D simulation selected from the top left, top right, bottom left, and middle of mixing space, see the markings A-D in Fig. 1 (panels with green background). In each panel, the top row of triangular heatmaps displays the concentration of pelagic algae (a), concentration of dissolved phosphorus (b), and pelagic algal specific growth rate (c) respectively. The vertical left edge in sub-panels (a)-(c) corresponds to the center of the lake, and the x-axis is the relative distance from the lake center. The compensation depth is marked in (a)-(c) with a horizontal dotted line. Sub-panel (d) in the bottom center displays the sedimented phosphorus surface density (black) and the dissolved phosphorus concentration along the bottom (blue) Sub-panel (e) in the bottom right displays the surface density of benthic algae where the color of the line indicates if the benthic algal growth is light-limited (blue) or nutrient-limited (red). The compensation depth is marked in (d)-(e) with a vertical dotted line. Note that the scales of the y-axes and color bars can differ between panels A-D. The table of values in the bottom left of each panel A-D lists the percentage of total phosphorus in each state variable.

#### 4.5.2 High horizontal and vertical mixing rates

Under well-mixed conditions (Figs. 7A, 8A, 9A), no spatial gradients in pelagic biomass and dissolved nutrients can arise (sub-panels a and b), but horizontal gradients in benthic biomass and sediment nutrients occur if benthic algal growth is light limited in at least some parts of the lake bottom (sub-panels d and e). In shallow, nutrient-poor pattern 1 lakes, both pelagic and benthic algal growth is nutrient limited in all but the deepest parts (Fig. 7Ac, e). Since there are no gradients in dissolved nutrient concentration, benthic biomass is constant down to the depth where light limitation kicks in (Fig. 7Ae). The stock of sediment nutrients is high and near evenly distributed across the lake bottom, since there are no spatial gradients in pelagic sinking losses (Fig. 7Ad). In more nutrient-rich and/or deeper pattern 2 lakes, large parts of the lake volume and bottom are aphotic and dissolved nutrient concentration is rather high. Pelagic and benthic algal growth is therefore restricted to a narrow band near the water surface and the shoreline, respectively (Fig. 8Ac, e). Consequently, benthic algae are absent from most of the lake bottom but reach a very high biomass peak closest to the shore, which is accompanied by a similar nearshore peak of sediment nutrients (Fig. 8Ad, e). In very deep and/or highly turbid pattern 3 lakes, light limitation drives pelagic algae extinct under well-mixed conditions. Consequently, no spatial gradients in dissolved nutrients can arise (Fig. 9Ab), while benthic algae survive in the shallow lake margins where they sustain a narrow band of sediment (Fig. 9Ad, e).

#### 4.5.3 Low horizontal and high vertical mixing

When horizontal mixing is low (corresponding to large lakes) but vertical mixing is high, sharp horizontal gradients develop in all state variables (Figs. 7B, 8B). Remarkably, the directions of these gradients differ between pattern 1 vs. pattern 2 and pattern 3 lakes. In shallow, nutrient-poor pattern 1 lakes, algal growth continues to be primarily nutrient limited (Fig. 7Be). Yet, because horizontal transport is limited, a large fraction of a lake’s nutrients can be ‘trapped’ in benthic algal biomass and sediment closest to the shore (Fig. 7Bd, e) where it is recycled locally, thus maintaining the highest nutrient concentrations, benthic and pelagic algal growth rates, and benthic biomass closest to the shore (Fig. 7Bb, c, e). Because sinking losses are disproportionally high at the shallowest depths, pelagic algae show the reverse pattern with lowest biomass in shallow nearshore areas (Fig. 7Ba). In more nutrient-rich and/or deeper pattern 2 and 3 lakes, algal growth is again restricted to a band near the water surface and the shoreline (Fig. 8Bc, e, 9Bc, e). The effects of limited horizontal transport are, however, opposite to pattern 1 lakes. Specifically, because of greater lake depth and/or stronger shading from pelagic algae, benthic algae are suppressed and restricted to shallow nearshore areas (Fig. 8Be, 9Be), while a large fraction of the system’s nutrients is maintained in dissolved form in central parts of the lake (Fig. 8Bb, 9Bb). Pelagic algae and dissolved nutrients therefore show the opposite horizontal distribution compared to pattern 1 lakes, with the lowest biomass and highest nutrient concentration in the lake center, where vertical mixing to aphotic depths restricts pelagic growth and nutrient uptake (Fig. 8Ba-c, 9Ba-c). In very deep and/or highly turbid pattern 3 lakes most of the lake volume is aphotic. Low horizontal mixing is therefore essential for the survival of pelagic algae because it allows for the buildup of pelagic algal biomass in nearshore areas with sufficient light.

In very deep and/or highly turbid pattern 3 lakes, low horizontal mixing uncouples shallow and deep parts of the lake, allowing pelagic algae to persist in nearshore areas where light is sufficient.

#### 4.5.4 Low vertical mixing

When vertical mixing is low, sharp vertical but no horizontal gradients arise in the pelagic state variables, regardless of the horizontal mixing rate. We therefore only show the case where horizontal mixing is low. Under these conditions, downward sinking of pelagic algae is much faster than the upward transport of dissolved nutrients.

Dissolved nutrients therefore accumulate in deeper parts of the lake but are scarce near the surface and the shoreline (Fig. 7Cb, 8Cb, 9Cb), and both pelagic and benthic production peak at a depth where light and nutrient supply are most balanced. This depth is near the bottom of the lake center in pattern 1 lakes and either near or at the lake surface in pattern 2 and 3 lakes, respectively (sub-panels c and e in Fig. 7C, 8C, 9C). In shallow, nutrient-poor pattern 1 lakes, the combination of low growth and excessive sinking losses drives pelagic algae to near-extinction (Fig. 7Ca). Benthic algae thus benefit from reduced competition, and the bulk of the system’s nutrients is approximately evenly distributed among benthic algae and the sediment, with a peak at the depth where algal growth switches from nutrient to light limitation (Fig. 7Cd, 7Ce).

In more nutrient-rich and/or deeper pattern 2 lakes, pelagic algal growth and sinking are more balanced. Consequently, pelagic algae can maintain a sizable population near the lake surface (Fig. 8Ca). Yet, compared to the low horizontal-high vertical mixing scenario, pelagic biomass is reduced and an even larger fraction of the system’s nutrients remains unused in dissolved form at aphotic depths (compare sub-panels a and b in Fig. 8B and 8C), while benthic algae survive at low biomass in the shallow margin of the lake (Fig. 8Ce). In very deep and/or highly turbid pattern 3 lakes, low vertical mixing prevents pelagic algae from being mixed to aphotic depths and allows them to maintain a sizeable population near the surface which shades out benthic algae (compare sub-panels Ce with Be and Ae in Fig. 9).

#### 4.5.5 Intermediate mixing

Finally, when both vertical and horizontal mixing rates are intermediate, all state variables exhibit spatial gradients that are dampened, but recognizable, combinations of the mixing scenarios described in the previous two sections. In shallow, nutrient-poor pattern 1 lakes, this results in a bimodal distribution of benthic algae and sediment nutrients, with one peak closest to the shoreline and another peak at the depth where algal growth switches from nutrient to light limitation (compare sub-panels d and e across Fig. 7B-D). Similarly, in all lake types, pelagic algae and dissolved nutrients show qualitatively similar counter-gradients as in the low horizontal mixing scenario and similar vertical gradients as in the low vertical mixing scenario (compare sub-panels a and b across Fig. 7B-D, 8B-D, 9B-D).

#### 4.5.6 Diagonal biomass patterns in mixing space

The effects of mixing on algal biomass described above can be summarized as follows: In pattern 1 lakes, low horizontal and vertical mixing is most beneficial to benthic but detrimental to pelagic algae, because pelagic algae experience the highest sinking losses under such conditions, liberating growth-limiting nutrients and minimizing shading to the benefit of benthic algae. In contrast, in pattern 2 lakes, high mixing is most beneficial to both benthic and pelagic algae because in such lakes growth is restricted to shallow depths and thus benefits from the nutrient supply to shallow areas provided by high mixing. Finally, in pattern 3 lakes, high horizontal and vertical mixing is most beneficial to benthic but detrimental to pelagic algae, because pelagic algae experience strong mixing to aphotic depths under such conditions, liberating growth-limiting nutrients and minimizing shading to the benefit of benthic algae. Moving from shallow, clear pattern 1 lakes via deeper and/or nutrient-rich pattern 2 lakes to very deep and/or turbid pattern 3 lakes, we therefore observe gradual shifts between opposite, diagonal biomass trends in mixing space for both benthic and pelagic algae. Specifically, in shallow, nutrient-poor pattern 1 lakes, benthic biomass is highest at low and lowest at high overall mixing, but shows the opposite trend in pattern 2 and pattern 3 lakes; conversely, pelagic biomass is highest at high and lowest at low overall mixing in pattern 1 and pattern 2 lakes but shows the opposite trend in pattern 3 lakes (Fig. 2, 7–9).

## 5 Discussion

The goal of the present study was to investigate how lake morphometry, nutrient content, water turbidity, and mixing intensity impacts algal dynamics. Our exploration across a comprehensive range of these environmental drivers using the Lake2D model revealed the following insights:

### 5.1 Analysis of algal patterns 1-3

Algal dynamics across the explored 5-dimensional parameter space can be categorized into three qualitatively distinct patterns, based on the contribution of pelagic vs. benthic algae to total lake biomass (patterns 1-3 in Fig. 2). In agreement with prior studies using 1D models, the shallow pattern 1 lakes exhibit low pelagic biomass due to high sinking losses [6, 48] Lake2D therefore predicts that pattern 1 would significantly diminish or disappear entirely if pelagic algae are neutrally boyant and/or can regulate their buoyancy (e.g., cyanobacteria [49–51]). If the nutrient content is increased, algal dynamics shift to the pattern 2 behavior due to increased pelagic algal growth that compensates for sinking losses. Conversely, if the mean depth is increased, pelagic algal sinking losses decrease, resulting in a shift to pattern 2 systems. Please note that the transition from one pattern to another is a continuous, smooth process with changing *N*_area_ and *z*_m_.

In pattern 2 lakes, sufficient overall light conditions ensure that high vertical mixing does not negatively impact pelagic algae. This leads to relatively small changes in depth-integrated biomass across mixing space, particularly for pelagic algal biomass, as evidenced by the lack of steep horizontal gradients in the pattern 2 panels of Fig. 6. Nutrient content, however, had the largest impact on total biomass due to increased pelagic algal biomass, indicating significant nutrient limitation in pelagic algal growth (blue A panels in Fig 4). Conversely, benthic algal biomass in pattern 2 lakes showed negligible increase with increasing *N*_area_ (white B panels in Fig. 4). We attribute this to two opposing effects: increased shading by pelagic algae limits benthic growth, while increased nutrient content promotes higher benthic algal densities in shallow areas. These compensatory effects result in relatively unchanged benthic algal biomass at the whole-lake scale as *N*_area_ is increased.

Our findings for the optically deeper pattern 3 lakes reveal an optimal vertical turbulence level that maximizes pelagic biomass, consistent with previous work [6, 52]. This phenomenon arises because sufficient vertical mixing both delivers adequate nutrients to the photic zone and counteracts algal sinking, thereby allowing pelagic algae to maintain high biomass. Conversely, pelagic algae either sink out of the photic zone if vertical mixing is too low or are mixed out of it if vertical mixing is too high.

Furthermore, Lake2D reveals a novel phenomenon for pattern 3 lakes: when the horizontal mixing rate is limited, pelagic algae persist even at high vertical mixing rates. Specifically, their survival is observed across all but the lowest vertical mixing rates (vertical bands in the A panels in Fig. 2 for *z*_m_ = 30 m). This phenomenon, which is only possible in 2D or 3D models and is thus overlooked by traditional 1D approaches, results from the interplay of two factors: horizontal mixing facilitates (1) the upward transport of dissolved nutrients from deep, poorly lit areas to shallow, well-lit areas, while (2) simultaneously being low enough to prevent excessive downward transport of pelagic algae to aphotic depths. Interestingly, when horizontal mixing is limited, pelagic algae can therefore sustain a sizable population even with high vertical mixing, since the slow horizontal transport allows pelagic algae to persist in shallow areas where light conditions are favorable regardless of the intensity of vertical mixing.

It is worth noting that Lake2D consistently predicts the extinction of pelagic algae under very low vertical mixing. However, this phenomenon primarily occurs at very low vertical mixing values that largely fall outside our depicted realistic simulation range, with the exception of pattern 1 lakes. Note also that for very low horizontal diffusive mixing (outside the depicted range in Fig. 2), whole-lake pelagic algal densities in pattern 3 lakes decline compared to the densities observed in the vertical bands. These very low horizontal mixing scenarios were omitted due to excessive computational runtimes.

### 5.2 Physical interpretation of mixing rates *d*_R_ **and** *d*_Z_

The mixing rates *d*_R_ and *d*_Z_ introduced in this study combine both diffusive mixing and lake radius/depth into single composite parameters. In general, combining parameters like this often enables efficient exploration of parameter space, where instead of varying each parameter independently one at a time, the composite parameter is varied instead, after which the underlying parameter values can then be inferred from the composite parameter. This approach reduces the total number of parameters that need to be varied to explore a parameter space. For example, the output of a simulation with *d*_R_ = 10 can be interpreted as any lake with *d*_r_ and *R* such that *d*_r_*/R*^2^ = 10.

However, in Lake2D, this approach can only be applied to the horizontal mixing rate *d*_R_ and not the vertical mixing rate *d*_Z_. This is because the pelagic sinking speed and the background light attenuation coefficient are independent of the mean depth, necessitating the inclusion of the mean depth as an independent variable when solving the non-dimensionalized model. This constraint does not apply to the horizontal mixing rate, implying that *d*_R_ can be varied as a single parameter (instead of varying *R* and *d*_z_ independently) without loss of generality.

Moreover, since the mean depth has to be treated as an independent variable, *d*_z_ can be calculated directly because *d*_Z_ is known. For lakes with a mean depth of 1 - 30 m, *d*_Z_ in the range 0.001 to 100 day^−1^ translates to *d*_z_ values in the range 10^−3^ 10^5^ m^2^ day^−1^. Comparing with empirically measured vertical diffusion coefficients in lakes, which span four orders of magnitude from 10^−1^ to 10^2^ m^2^ day^−1^ [53–55], demonstrates that our explored range of horizontal mixing comprehensively covers all plausible empirical values.

While the vertical diffusion coefficients can be calculated directly from the vertical mixing rates, calculating horizontal diffusion coefficients from the horizontal mixing rates requires an assumption about the size of the lake area (this parameter is not present in the non-dimensionalized model). The size distribution of lakes on Earth is vast, spanning nine orders of magnitude (depending on the minimum lake size threshold) and ranges from a fraction of a hectare to tens of thousands of square kilometers for the very largest lakes [56]. If we consider lake areas from 10^4^ to 10^10^ m^2^ (*>*99% of all lakes on Earth [57]) and the *d*_Z_ values used in this study (10^−3^ to 10 day^−1^), these values correspond to horizontal diffusion coefficients in the range 1 to 10^10^ m^2^ day^−1^, which covers the entire empirically observed range of horizontal diffusion in lakes which spans from 10 to 10^8^ m^2^ day^−1^ [44, 54, 58, 59].

### 5.3 Benthic deep chlorophyll maximum

A deep chlorophyll maximum (DCM) refers to the phenomenon where algal biomass concentration peaks at some intermediate depth. Empirically, DCMs are readily observed in pelagic algae [60, 61]; however, measurements of benthic algal biomass across a depth transect are uncommon [9], and to our knowledge the formation of benthic algal DCMs due to posing gradients in light intensity and dissolved nutrients has not been proposed. However, models predict intermediate peaks in benthic production [25, 43, 62–64], indicating the possibility of the formation of benthic algal DCMs in lakes.

Interestingly, Lake2D predicts the formation of a benthic DCM in shallow lakes (*z*_m_ ≤ 3m) with low vertical mixing (see Fig. 7Ce-De), where benthic areal biomass peaks at an intermediate depth as growth switches from nutrient limitation to light limitation (transition from blue to red in 7Ce-De). For a benthic DCM to form, three conditions must be met: (1) light intensity must decrease appreciably with depth; (2) dissolved P concentration must increase with depth; (3) benthic algal growth must switch from light limitation in shallow areas to nutrient limitation in deeper areas at some intermediate depth. When these three conditions are met, a benthic DCM will form at the depth where this switch occurs.

Condition (1) requires a combination of sufficient depth and light attenuation per unit depth for a non-negligible vertical gradient in light intensity to form, and is met in all but exceptionally clear and shallow lakes. For example, in a lake that is 1 m deep with a background turbidity *k*_bg_ = 0.2, light intensity has decreased by 18% by the time it reaches the bottom (not accounting for shading by pelagic algae).

Condition (2) is met only when vertical mixing is low enough to allow vertical gradients to form in the pelagic. Moreover, there must also be some process present that transports nutrients into the deeper areas. If not, turbulent mixing will eventually erase any gradient in dissolved P, no matter how low the vertical mixing rate is. In lake2D, this role is filled by sinking pelagic algae, but there are alternative mechanisms such as the resuspension and sinking of sedimented detritus or benthic algae. Therefore, a consequence of the extinction of pelagic algae in Lake2D is a well-mixed pelagic zone at steady state, rendering a benthic DCM impossible in such scenarios.

Finally, condition (3) is met when dissolved P concentrations are low enough towards the surface of the lake to cause benthic algal growth to be light-limited there, and high enough in deeper areas such that benthic algal growth is light-limited at greater depths.

In our simulations, these conditions are only realized when the vertical mixing is sufficiently low to allow vertical gradients in dissolved P (condition (2)), and condition (1) is met in all explored scenarios. If the water is turbid (high *k*_bg_), and/or nutrient content is high (resulting in significant shading from pelagic algae), then condition (3) is not met and benthic algal growth is light-limited with the highest biomass concentrations in the shallowest areas. We therefore do not observe benthic DCMs for *N*_area_ = 3000 mg P m^−3^, since this high nutrient content causes too much shading from pelagic algae and an abundance of dissolved P.

In the literature, wave action [62, 64, 65] and ice scouring [10, 66, 67] (abrasive erosion along lake margins from surface waves and moving ice) are commonly cited as causes of a decrease in benthic algal densities in shallow areas, which together with reduced algal growth in deeper areas due to light-limitation would lead to the formation of benthic algal biomass peaks at intermediate water depths. While wave action and ice scouring most likely contribute to a decrease in benthic algal densities in shallow lake margins, our simulations show that benthic DCMs can form independently of these processes. Yet, including mechanical scour and resuspension of both benthic algae and sediment would be an obvious next step in the development of model Lake2D.

## Appendix

### A Non-dimensionalized system

The non-dimensionalized system is given by

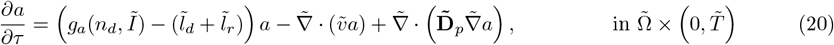

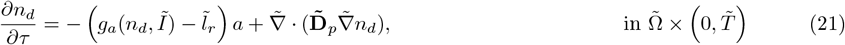

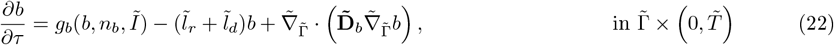

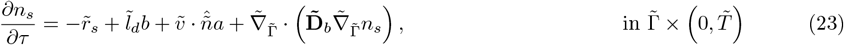

where 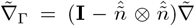 denotes the tangential gradient along the lake bottom 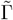, and 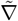 is the usual gradient operator in the non-dimensionalized space. Lowercase letters and tildes are systematically used to indicate the non-dimensionalized forms of the original parameters, state variables, vectors, and operators. The forms *I, g*_*a*_, and *g*_*b*_ are given by

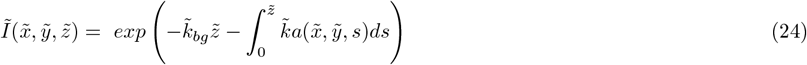

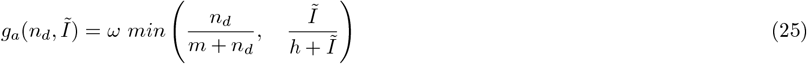

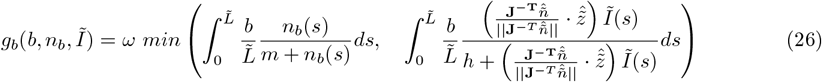

with lake-surface boundary conditions

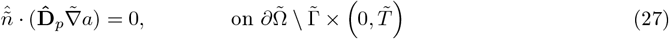

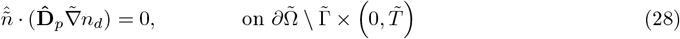

along the lake surface and

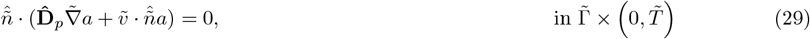

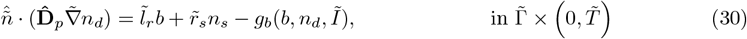

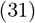

along the lake bottom. The non-dimensionalized equation for the dissolved nutrients inside the benthic algal layer is given by

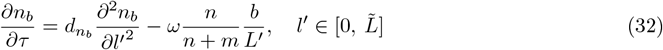

With the boundary condition

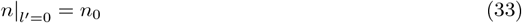

on the surface of the benthic algal layer and

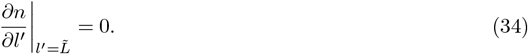

on the bottom of the benthic algal layer. Non-dimensionalized variables and parameters are given by

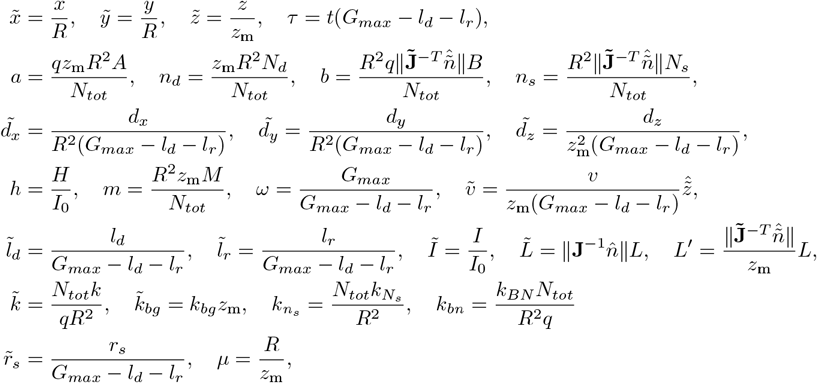

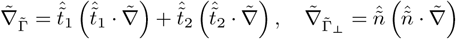, where 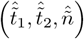 is a basis on the lake bottom, where 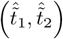 spans the bottom surface and 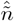 is the lake bottom normal vector.

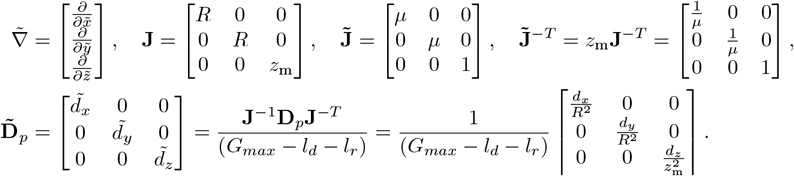

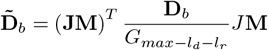, where 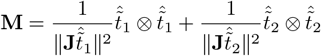, the operator ⊗ is the outer product defined as 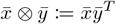 for two arbitrary column vectors 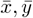.

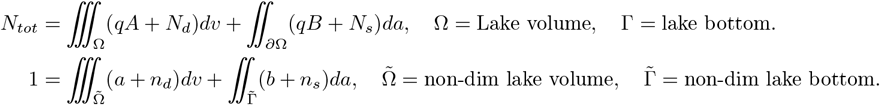

## Supplementary data with varying background turbidity

**Figure B1:**
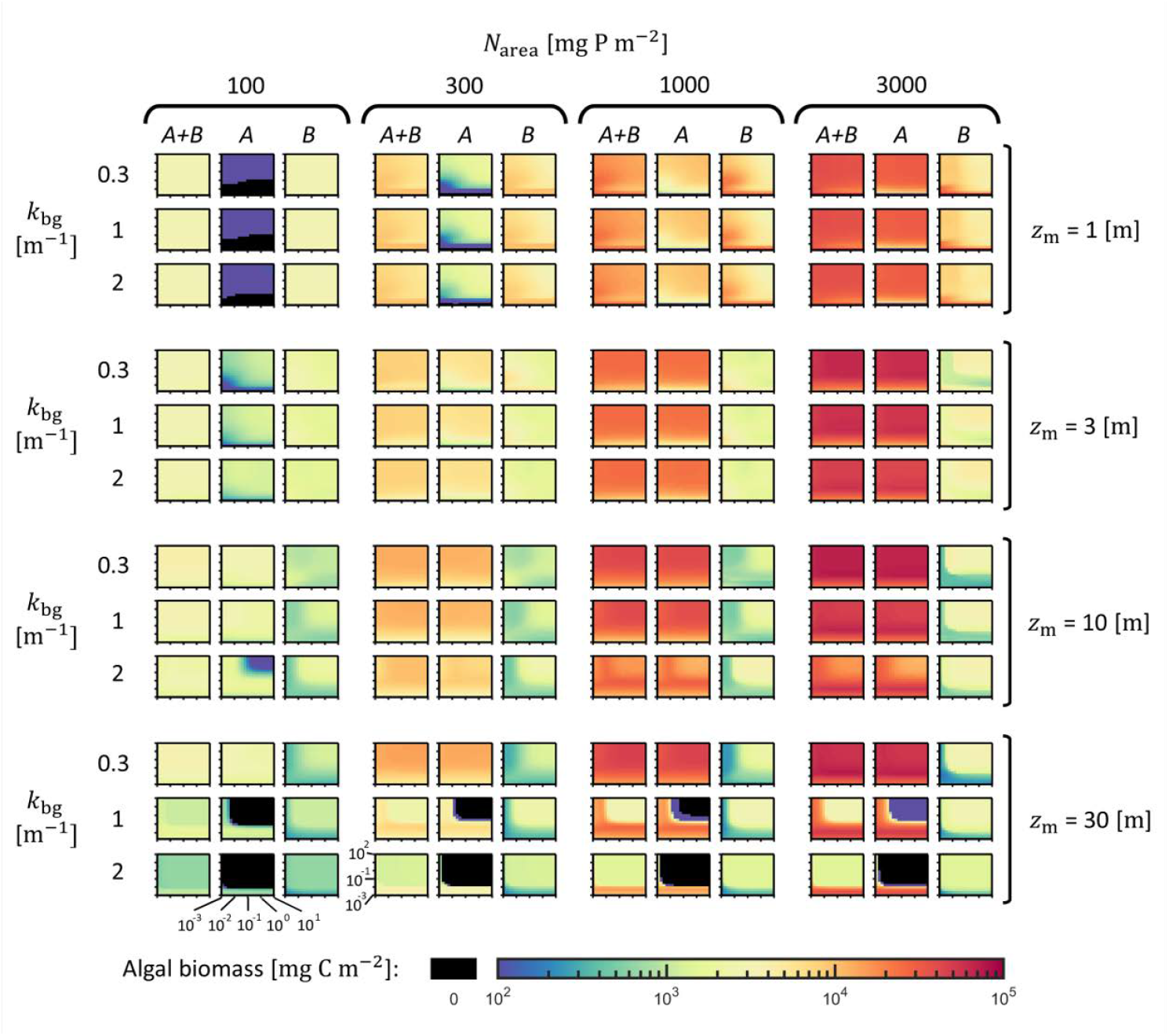
Heatmaps of lake-wide pelagic (A), benthic (B), and total (A+B) algal biomass for varying total nutrient content per surface area N_tot_, mean depth z_m_, and diffusive mixing rates d_R_ and d_Z_, and background turbidity k_bg_. The horizontal mixing rate d_R_ varies along the horizontal axis of each panel in the range [0.001, 10] day^−1^, and the vertical mixing rate d_Z_ varies along the vertical axis of each panel in the range [0.001, 100] day^−1^. Both axes are plotted on a log10 scale. Each pixel in a panel is the average biomass per unit of lake surface area (mg C m^−2^) on a log10 scale (see color bar at the bottom of the figure). Black areas indicate extinction, defined as an average concentration < 0.001 mg C m^−3^.

## References

[1] E. Fee, R. Hecky, S. Kasian, and D. Cruikshank, “Effects of lake size, water clarity, and climatic variability on mixing depths in Canadian Shield lakes,” Limnology and oceanography, vol. 41, no. 5, pp. 912–920, 1996.

[2] B. Qin, J. Zhou, J. J. Elser, W. S. Gardner, J. Deng, and J. D. Brookes, “Water depth underpins the relative roles and fates of nitrogen and phosphorus in lakes,” Environmental Science & Technology, vol. 54, no. 6, pp. 3191–3198, 2020.

[3] L. Zhao, R. Zhu, Q. Zhou, E. Jeppesen, and K. Yang, “Trophic status and lake depth play important roles in determining the nutrient-chlorophyll a relationship: Evidence from thousands of lakes globally,” Water Research, vol. 242, p. 120182, 2023.

[4] C. Morana, A. V. Borges, L. Deirmendjian, W. Okello, H. Sarmento, J.-P. Descy, I. A. Kimirei, and S. Bouillon, “Prevalence of autotrophy in non-humic African lakes,” Ecosystems, vol. 26, no. 3, pp. 627–642, 2023.

[5] Y. Vadeboncoeur, G. Peterson, M. J. Vander Zanden, and J. Kalff, “Benthic algal production across lake size gradients: interactions among morphometry, nutrients, and light,” Ecology, vol. 89, no. 9, pp. 2542–2552, 2008.

[6] C. G. Jäger, S. Diehl, and M. Emans, “Physical determinants of phytoplankton production, algal stoichiometry, and vertical nutrient fluxes,” The American Naturalist, vol. 175, no. 4, pp. E91–E104, 2010.

[7] L. Håkanson, “The importance of lake morphometry for the structure and function of lakes,” International Review of Hydrobiology: A Journal Covering all Aspects of Limnology and Marine Biology, vol. 90, no. 4, pp. 433–461, 2005.

[8] S. Brothers, Y. Vadeboncoeur, and P. Sibley, “Benthic algae compensate for phytoplankton losses in large aquatic ecosystems,” Global Change Biology, vol. 22, no. 12, pp. 3865–3873, 2016.

[9] I. Puts, A.-K. Bergström, H. Verheijen, S. Norman, and J. Ask, “An ecological and methodological assessment of benthic gross primary production in northern lakes,” Ecosphere, vol. 13, no. 3, p. e3973, 2022.

[10] D. Seekell and B. Cael, “Why does the relationship between benthic primary production and lake morphometry vary regionally?,” Aquatic Sciences, vol. 85, no. 3, p. 87, 2023.

[11] B. Althouse, S. Higgins, and M. J. Vander Zanden, “Benthic and planktonic primary production along a nutrient gradient in Green Bay, Lake Michigan, USA,” Freshwater Science, vol. 33, no. 2, pp. 487–498, 2014.

[12] A. Kuczynski, M. T. Auer, C. N. Brooks, and A. G. Grimm, “The Cladophora resurgence in Lake Ontario: characterization and implications for management,” Canadian journal of fisheries and aquatic sciences, vol. 73, no. 6, pp. 999–1013, 2016.

[13] F. Cremona, A. Laas, L. Arvola, D. Pierson, P. Nõges, and T. Nõges, “Numerical exploration of the planktonic to benthic primary production ratios in lakes of the Baltic Sea catchment,” Ecosystems, vol. 19, pp. 1386–1400, 2016.

[14] A. R. McCormick, J. S. Phillips, J. C. Botsch, and A. R. Ives, “Shifts in the partitioning of benthic and pelagic primary production within and across summers in Lake My`vatn, Iceland,” Inland Waters, vol. 11, no. 1, pp. 13–28, 2021.

[15] D. Krause-Jensen, S. Markager, and T. Dalsgaard, “Benthic and pelagic primary production in different nutrient regimes,” Estuaries and coasts, vol. 35, pp. 527–545, 2012.

[16] L. Soares and M. Calijuri, “Deterministic modelling of freshwater lakes and reservoirs: Current trends and recent progress,” Environmental Modelling and Software, vol. 144, p. 105143, 2021.

[17] J. Huisman, P. van Oostveen, and F. J. Weissing, “Critical depth and critical turbulence: two different mechanisms for the development of phytoplankton blooms,” Limnology and oceanography, vol. 44, no. 7, pp. 1781–1787, 1999.

[18] M. Scheffer, “Multiplicity of stable states in freshwater systems,” Hydrobiologia, vol. 200, no. 1, pp. 475–486, 1990.

[19] A. B. Janssen, G. B. Arhonditsis, A. Beusen, K. Bolding, L. Bruce, J. Bruggeman, R.-M. Couture, A. S. Downing, J. Alex Elliott, M. A. Frassl, et al., “Exploring, exploiting and evolving diversity of aquatic ecosystem models: a community perspective,” Aquatic ecology, vol. 49, pp. 513–548, 2015.

[20] D. M. Imboden, “Phosphorus model of lake eutrophication,” Limnology and Oceanography, vol. 19, no. 2, pp. 297–304, 1974.

[21] C. R. Fragoso Jr, E. H. van Nes, J. H. Janse, and D. da Motta Marques, “IPH-TRIM3D-PCLake: A three-dimensional complex dynamic model for subtropical aquatic ecosystems,” Environmental Modelling & Software, vol. 24, no. 11, pp. 1347–1348, 2009.

[22] F. Hu, K. Bolding, J. Bruggeman, E. Jeppesen, M. R. Flindt, L. Van Gerven, J. H. Janse, A. B. Janssen, J. J. Kuiper, W. M. Mooij, et al., “FABM-PCLake - linking aquatic ecology with hydrodynamics,” Geoscientific Model Development, vol. 9, no. 6, pp. 2271–2278, 2016.

[23] E. J. Fee, “A relation between lake morphometry and primary productivity and its use in interpreting whole-lake eutrophication experiments,” Limnology and Oceanography, vol. 24, no. 3, pp. 401–416, 1979.

[24] S. R. Carpenter, “Lake geometry: implications for production and sediment accretion rates,” Journal of Theoretical Biology, vol. 105, no. 2, pp. 273–286, 1983.

[25] S. P. Devlin, M. J. Vander Zanden, and Y. Vadeboncoeur, “Littoral-benthic primary production estimates: Sensitivity to simplifications with respect to periphyton productivity and basin morphometry,” Limnology and Oceanography: Methods, vol. 14, no. 2, pp. 138–149, 2016.

[26] T. K. Andersen, K. Bolding, A. Nielsen, J. Bruggeman, E. Jeppesen, and D. Trolle, “How morphology shapes the parameter sensitivity of lake ecosystem models,” Environmental Modelling & Software, vol. 136, p. 104945, 2021.

[27] R. Sachse, T. Petzoldt, M. Blumstock, S. Moreira, M. Pätzig, J. Rücker, J. H. Janse, W. M. Mooij, and S. Hilt, “Extending one-dimensional models for deep lakes to simulate the impact of submerged macrophytes on water quality,” Environmental Modelling & Software, vol. 61, pp. 410–423, 2014.

[28] I. A. Oleksy, C. T. Solomon, S. E. Jones, C. Olson, B. L. Bertolet, R. Adrian, S. Bansal, J. S. Baron, S. Brothers, S. Chandra, et al., “Controls on lake pelagic primary productivity: Formalizing the nutrient-color paradigm,” Journal of Geophysical Research: Biogeosciences, vol. 129, no. 12, p. e2024JG008140, 2024.

[29] F. Rivera Vasconcelos, S. Diehl, P. Rodríguez, J. Karlsson, and P. Byström, “Effects of terrestrial organic matter on aquatic primary production as mediated by pelagic–benthic resource fluxes,” Ecosystems, vol. 21, pp. 1255–1268, 2018.

[30] M. Genkai-Kato and S. R. Carpenter, “Eutrophication due to phosphorus recycling in relation to lake morphometry, temperature, and macrophytes,” Ecology, vol. 86, no. 1, pp. 210–219, 2005.

[31] P. C. Hanson, A. I. Pollard, D. L. Bade, K. Predick, S. R. Carpenter, and J. A. Foley, “A model of carbon evasion and sedimentation in temperate lakes,” Global Change Biology, vol. 10, no. 8, pp. 1285–1298, 2004.

[32] P. T. Kelly, C. T. Solomon, J. A. Zwart, and S. E. Jones, “A framework for understanding variation in pelagic gross primary production of lake ecosystems,” Ecosystems, vol. 21, pp. 1364–1376, 2018.

[33] Z. Tan, Q. Zhuang, N. J. Shurpali, M. E. Marushchak, C. Biasi, W. Eugster, and K. Walter Anthony, “Modeling CO_2_ emissions from Arctic lakes: Model development and site-level study,” Journal of Advances in Modeling Earth Systems, vol. 9, no. 5, pp. 2190–2213, 2017.

[34] M. Genkai-Kato, Y. Vadeboncoeur, L. Liboriussen, and E. Jeppesen, “Benthic–planktonic coupling, regime shifts, and whole-lake primary production in shallow lakes,” Ecology, vol. 93, no. 3, pp. 619– 631, 2012.

[35] C. F. Cerco and T. Cole, “Three-dimensional eutrophication model of Chesapeake Bay,” Journal of Environmental Engineering, vol. 119, no. 6, pp. 1006–1025, 1993.

[36] C. R. Fragoso Jr, D. M. M. Marques, W. Collischonn, C. E. Tucci, and E. H. Van Nes, “Modelling spatial heterogeneity of phytoplankton in Lake Mangueira, a large shallow subtropical lake in South Brazil,” Ecological modelling, vol. 219, no. 1-2, pp. 125–137, 2008.

[37] J. Skerratt, K. Wild-Allen, F. Rizwi, J. Whitehead, and C. Coughanowr, “Use of a high resolution 3d fully coupled hydrodynamic, sediment and biogeochemical model to understand estuarine nutrient dynamics under various water quality scenarios,” Ocean & coastal management, vol. 83, pp. 52–66, 2013.

[38] F. Soulignac, P.-A. Danis, D. Bouffard, V. Chanudet, E. Dambrine, Y. Guénand, T. Harmel, B. W. Ibelings, D. Trevisan, R. Uittenbogaard, et al., “Using 3d modeling and remote sensing capabilities for a better understanding of spatio-temporal heterogeneities of phytoplankton abundance in large lakes,” Journal of Great Lakes Research, vol. 44, no. 4, pp. 756–764, 2018.

[39] H. Harlin, K. Larsson, and S. Diehl, “Can whole-lake algal biomass be captured by one-dimensional modeling approaches? An exploration using ‘Lake2D’,” bioRxiv, pp. 2025–05, 2025.

[40] J. Huisman and F. J. Weissing, “Light-limited growth and competition for light in well-mixed aquatic environments: an elementary model,” Ecology, vol. 75, no. 2, pp. 507–520, 1994.

[41] L. Håkanson, “On lake form, lake volume and lake hypsographic survey,” Geografiska Annaler: Series A, Physical Geography, vol. 59, no. 1-2, pp. 1–29, 1977.

[42] J. Stachelek, P. J. Hanly, and P. A. Soranno, “Imperfect slope measurements drive overestimation in a geometric cone model of lake and reservoir depth,” Inland Waters, vol. 12, no. 2, pp. 283–293, 2022.

[43] M. Klaus, H. A. Verheijen, J. Karlsson, and D. A. Seekell, “Depth and basin shape constrain ecosystem metabolism in lakes dominated by benthic primary producers,” Limnology and Oceanography, vol. 67, no. 12, pp. 2763–2778, 2022.

[44] F. Peeters, A. Wüest, G. Piepke, and D. M. Imboden, “Horizontal mixing in lakes,” Journal of Geophysical Research: Oceans, vol. 101, no. C8, pp. 18361–18375, 1996.

[45] B. Cael, A. Heathcote, and D. Seekell, “The volume and mean depth of Earth’s lakes,” Geophysical Research Letters, vol. 44, no. 1, pp. 209–218, 2017.

[46] M. Chen, G. Zeng, J. Zhang, P. Xu, A. Chen, and L. Lu, “Global landscape of total organic carbon, nitrogen and phosphorus in lake water,” Scientific reports, vol. 5, no. 1, p. 15043, 2015.

[47] S. Sobek, L. J. Tranvik, Y. T. Prairie, P. Kortelainen, and J. J. Cole, “Patterns and regulation of dissolved organic carbon: An analysis of 7,500 widely distributed lakes,” Limnology and oceanography, vol. 52, no. 3, pp. 1208–1219, 2007.

[48] S. Diehl, “Phytoplankton, light, and nutrients in a gradient of mixing depths: theory,” Ecology, vol. 83, no. 2, pp. 386–398, 2002.

[49] G. Fogg and A. Walsby, “Buoyancy regulation and the growth of planktonic blue-green algae,” Internationale Vereinigung für Theoretische und Angewandte Limnologie: Mitteilungen, vol. 19, no. 1, pp. 182–188, 1971.

[50] R. Thomas and A. Walsby, “Buoyancy regulation in a strain of microcystis,” Microbiology, vol. 131, no. 4, pp. 799–809, 1985.

[51] A. E. Walsby, P. K. Hayes, R. Boje, and L. J. Stal, “The selective advantage of buoyancy provided by gas vesicles for planktonic cyanobacteria in the Baltic Sea,” The New Phytologist, vol. 136, no. 3, pp. 407–417, 1997.

[52] J. Huisman, M. Arrayás, U. Ebert, and B. Sommeijer, “How do sinking phytoplankton species manage to persist?,” The American Naturalist, vol. 159, no. 3, pp. 245–254, 2002.

[53] J. Huisman, J. Sharples, J. M. Stroom, P. M. Visser, W. E. A. Kardinaal, J. M. Verspagen, and B. Sommeijer, “Changes in turbulent mixing shift competition for light between phytoplankton species,” Ecology, vol. 85, no. 11, pp. 2960–2970, 2004.

[54] D. Imboden and S. Emerson, “Natural radon and phosphorus as limnologic tracers: Horizontal and vertical eddy diffusion in Greifensee,” Limnology and Oceanography, vol. 23, no. 1, pp. 77–90, 1978.

[55] B. Boehrer, J. Ilmberger, and K. O. Münnich, “Vertical structure of currents in western Lake Constance,” Journal of Geophysical Research: Oceans, vol. 105, no. C12, pp. 28823–28835, 2000.

[56] B. B. Cael and D. A. Seekell, “The size-distribution of earth’s lakes,” Scientific reports, vol. 6, no. 1, p. 29633, 2016.

[57] C. Verpoorter, T. Kutser, D. A. Seekell, and L. J. Tranvik, “A global inventory of lakes based on high-resolution satellite imagery,” Geophysical Research Letters, vol. 41, no. 18, pp. 6396–6402, 2014.

[58] F. Peeters and H. Hofmann, “Length-scale dependence of horizontal dispersion in the surface water of lakes,” Limnology and Oceanography, vol. 60, no. 6, pp. 1917–1934, 2015.

[59] C. Murthy, “Horizontal diffusion characteristics in Lake Ontario,” Journal of physical oceanography, vol. 6, no. 1, pp. 76–84, 1976.

[60] A. E. Scofield, J. M. Watkins, E. Osantowski, and L. G. Rudstam, “Deep chlorophyll maxima across a trophic state gradient: A case study in the Laurentian Great Lakes,” Limnology and Oceanography, vol. 65, no. 10, pp. 2460–2484, 2020.

[61] T. H. Leach, B. E. Beisner, C. C. Carey, P. Pernica, K. C. Rose, Y. Huot, J. A. Brentrup, I. Domaizon, H.-P. Grossart, B. W. Ibelings, et al., “Patterns and drivers of deep chlorophyll maxima structure in 100 lakes: The relative importance of light and thermal stratification,” Limnology and Oceanography, vol. 63, no. 2, pp. 628–646, 2018.

[62] R. J. Stevenson and E. F. Stoermer, “Quantitative differences between benthic algal communities along a depth gradient in Lake Michigan,” Journal of Phycology, vol. 17, p. 29–36, Mar. 1981.

[63] C. A. Gushulak, H. A. Haig, M. V. Kingsbury, B. Wissel, B. F. Cumming, and P. R. Leavitt, “Effects of spatial variation in benthic phototrophs along a depth gradient on assessments of whole-lake processes,” Freshwater Biology, vol. 66, no. 11, pp. 2118–2132, 2021.

[64] Y. Vadeboncoeur, S. P. Devlin, P. B. McIntyre, and M. J. Vander Zanden, “Is there light after depth? distribution of periphyton chlorophyll and productivity in lake littoral zones,” Freshwater science, vol. 33, no. 2, pp. 524–536, 2014.

[65] K. D. Hoagland and C. G. Peterson, “Effects of light and wave disturbance on vertical zonation of attached microalgage in a large reservoir,” Journal of Phycology, vol. 26, no. 3, pp. 450–457, 1990.

[66] M. Page, T. Goldhammer, S. Hilt, S. Tolentino, and S. Brothers, “Filamentous algae blooms in a large, clear-water lake: Potential drivers and reduced benthic primary production,” Water, vol. 14, no. 13, p. 2136, 2022.

[67] P. N. Bégin, M. Rautio, Y. Tanabe, M. Uchida, A. I. Culley, and W. F. Vincent, “The littoral zone of polar lakes: inshore–offshore contrasts in an ice-covered High Arctic lake,” Arctic Science, vol. 7, no. 1, pp. 158–181, 2020.

